# Mild focal cooling selectively impacts computations in dendritic trees

**DOI:** 10.1101/2024.11.02.621672

**Authors:** Meisam Habibi Matin, Shulan Xiao, Krishna Jayant

**Author notes:** Correspondence should be addressed to Krishna Jayant.

## Abstract

Focal cooling is a powerful technique to temporally scale neural dynamics. However, the underlying cellular mechanisms causing this scaling remain unresolved. Here, using targeted focal cooling (with a spatial resolution of 100 micrometers), dual somato-dendritic patch clamp recordings, two-photon calcium imaging, transmitter uncaging, and modeling we reveal that a 5°C drop can enhance synaptic transmission, plasticity, and input-output transformations in the distal apical tuft, but not in the basal dendrites of intrinsically bursting L5 pyramidal neurons. This enhancement depends on N-methyl-D-aspartate (NMDA) and Kv4.2, suggesting electrical structure modulation. Paradoxically, and despite the increase in tuft excitability, we observe a reduced rate of recovery from inactivation for apical Na+ channels, thereby regulating back-propagating action potential invasion, coincidence detection, and overall burst probability, resulting in an “apparent” slowing of somatic spike output. Our findings reveal a differential temperature sensitivity along the basal-tuft axis of L5 neurons analog modulates cortical output.

## INTRODUCTION

Compared to neuromodulation techniques such as optogenetics and chemogenetics, which digitally control neural circuit activity by turning it on or off^1,2^, mild focal cooling of the brain allows for graded (analog) changes in temporal dynamics^3,4^. This temporal scaling has been found to influence behavior and reveal the circuit functions involved in timing-dependent computations. For example, studies have shown that cooling a specific brain region in zebra finches can elongate the birds’ song motifs while preserving their acoustic microstructure^5^. On the other hand, cooling the striatum affected timing judgment without influencing choice behavior^6^.

Applying cooling to the brain’s surface can change the temporal pattern of behavior without disrupting its structure^4^, but it’s still unclear how these focal temperature changes affect circuit dynamics. To address this question, this study uses a range of biophysical measures to elucidate the impact of temperature changes on layer-5 (L5) pyramidal neurons - the primary output layer of the cortex.

L5 pyramidal neurons mediate higher-order cognitive functions, such as motor control and sensory integration^7–9^, and their extensive axonal projections and complex synaptic interactions render them crucial in cortical information processing and connectivity^9–11^. Thus, understanding the biophysical basis underlying focal cooling-induced L5 modulation and its impact on excitability is pivotal for advancing our knowledge surrounding L5 cortical excitability and plasticity – a central focus in our study.

Computational models and experiments have shown that lowering temperature can affect fundamental characteristics of neuronal membranes, such as their ability to modulate excitability^12–15^, reduce membrane fluidity^16,17^, influence metabolism^18^, impact impulse propagation^13,19–24^, and regulate synaptic transmission^25–29^. However, how selective/focal temperature changes affect neural coding remains unclear, especially across different cellular domains. Such focal reduction in temperature effects could locally modulate the temperature dependence of voltage-gated ion channel activity, influence plasticity, and eventual integration in a compartmentalized fashion, selectively modulating neuronal signal processing – a feature we uncover in L5 neurons.

L5 pyramidal neurons have distinct dendritic structures—tufted dendrites in the upper cortical layers and basal dendrites in the deeper cortical layers — each compartmentalized for different types of synaptic integration^30–34^. Tuft dendrites integrate inputs from higher cortical areas, influencing complex cognitive functions such as attention and perception^7,35^. On the other hand, basal dendrites primarily integrate local cortical and subcortical inputs, contributing to sensory and motor information processing^9,36,37^. Understanding how temperature affects these different dendritic regions from an integration standpoint can reveal how thermal changes might influence the integration of diverse feedforward and feedback synaptic streams^38–41^, which could lead to significant effects downstream of the cortex – a prospect not farfetched when focal cooling is applied from the brain surface.

Temperature can also significantly impact synaptic plasticity mechanisms like long-term potentiation (LTP) and long-term depression (LTD)^42,43^, which are crucial for learning and memory^44^. Since tuft and basal dendrites have distinct synaptic and signaling pathways^11,45–49^, temperature changes could affect these processes differently, for example, via changes in calcium dynamics^50^ or local excitability^51^. Understanding how temperature influences plasticity in these distinct dendrites can give us insights into how thermal stress affects cognitive functions and adaptive learning processes. Additionally, the reversible nature of focal cooling would also allow us to observe how plasticity processes recover after the cooling is removed. These recordings can provide insights into the adaptability and resilience of neural circuits.

In this study, using focal cooling with a resolution of ∼100 microns, dual somato-dendritic patch clamp recordings, two-photon calcium imaging, and transmitter uncaging, we reveal that a temperature drop of only 5°C from a physiological baseline can enhance synaptic transmission, plasticity, and input-output transformations in the distal apical tuft, but not in the basal dendrites of intrinsically bursting L5 pyramidal neurons. Our recordings suggest that this amplification of excitability and plasticity is tied to both Kv4.2 and N-methyl-D-aspartate (NMDA) activity, revealing a gradient in cooling-induced circuit tuning along the apical dendritic axis and suggesting that moderate and mild focal cooling could be used as an additional knob to differentially fine-tune integration in the tuft while keeping the basal properties intact. This is specifically critical as previous studies are at odds with each other. For example, some reports have indicated that neuronal excitability increases in the temperature range of 15-20 °C before being suppressed around 10 °C^13,27^. Conversely, some studies show a net slowing of spike output as a function of temperature, although within a moderate fluctuation range (ΔT <10°C)^52^. Here, we illustrate that not all neuronal locations respond to temperature similarly, so the net output as a function of temperature represents a unique integration of mechanisms.

Our findings reveal that focal cooling can dramatically enhance synaptic transmission, excitability, and plasticity in the distal apical tuft of layer 5 pyramidal neurons with just a 5°C temperature drop, but not across basal dendrites. This differential effect, dependent on NMDA receptors and Kv4.2 channels, not only amplifies neural dynamics locally but also introduces a fascinating paradox. While boosting neuronal excitability locally, it simultaneously slows the recovery of Na+ channels from inactivation, subtly modulating spike output and reducing burst probability. These novel findings not only shed light on the complex biophysics of focal cooling-induced neuromodulation but also suggest new ways for manipulating neuronal coding and signal processing in an analog fashion, paving the way for therapeutic innovations.

## RESULTS

### Mild focal cooling amplifies tuft dendritic plasticity and excitability

To reveal how focal cooling modulates dendritic excitability and plasticity across the tuft and basal dendrites of L5 pyramidal neurons, we combined focal cooling using a custom-designed probe (**Fig. 1 and Fig. S1**) with whole-cell somatic patch clamp, field stimulation, and two-photon imaging (**Fig. 1**). Following a recent report in which a novel form of N-methyl-D-aspartate receptors (NMDARs) and Kv4.2-dependent potentiation in the tuft was uncovered, we stimulated tuft dendrites with low frequency (0.1 Hz) single unpaired EPSPs (see Methods) to induce plasticity^48^ (**Fig. 1A**). We performed this stimulation at a physiological baseline (35°C) and under mild focal cooling to a final temperature of 30°C, separately across different cells (**Fig. 1B**). It is critical to note that this temperature change was calibrated and the range over which cooling is effectively imparted was extracted and found to be ∼100µm (**Fig. S1)** and we further corroborated the design of the probe through thermal modeling of the tissue in the chamber (**Fig. S2A, B**). The initial pre-stimulation baseline and final post-plasticity readouts were performed at the physiological baseline temperature of 35^°^C and stimulation frequency of 1 Hz. It is critical to note that the plasticity induction and recordings were done under bath application of Gabazine (**Fig. S3, and see Methods**). This gave a stable EPSP amplitude to compare the effect of mild focal cooling on plasticity. We observed that the resultant amplitude change during plasticity induction was higher under mild focal cooling. The eventual post-induction readout at 35°C reflected an increased excitatory post-synaptic response after mild focal cooling compared to the case in which no cooling was performed. This result, shown for a single neuron here, suggests that mild focal cooling during tuft dendritic stimulation amplifies the responsivity, which persists long after focal cooling is removed (**Fig. S4**). This post-induction recording at 1Hz was performed at least 10 minutes after the low-frequency induction to ensure that the potentiation effect is not transient. It should be noted that a full recovery from 30°C to 35°C takes 3-4 minutes (**Fig. S1**), and the post-LTP recording was carried out after a 10-minute settling-down period. We observed that when LTP is induced under mild focal cooling, the EPSP amplitude increase as a function of field stimulation is significantly higher than when LTP is inducted at 35°C (49%±9.8% compared to 22%±6.4%, Wilcoxon Mann Whitney test, n=7 neurons, p=0.0346) (**Fig. 1C**).

**Fig. 1.**
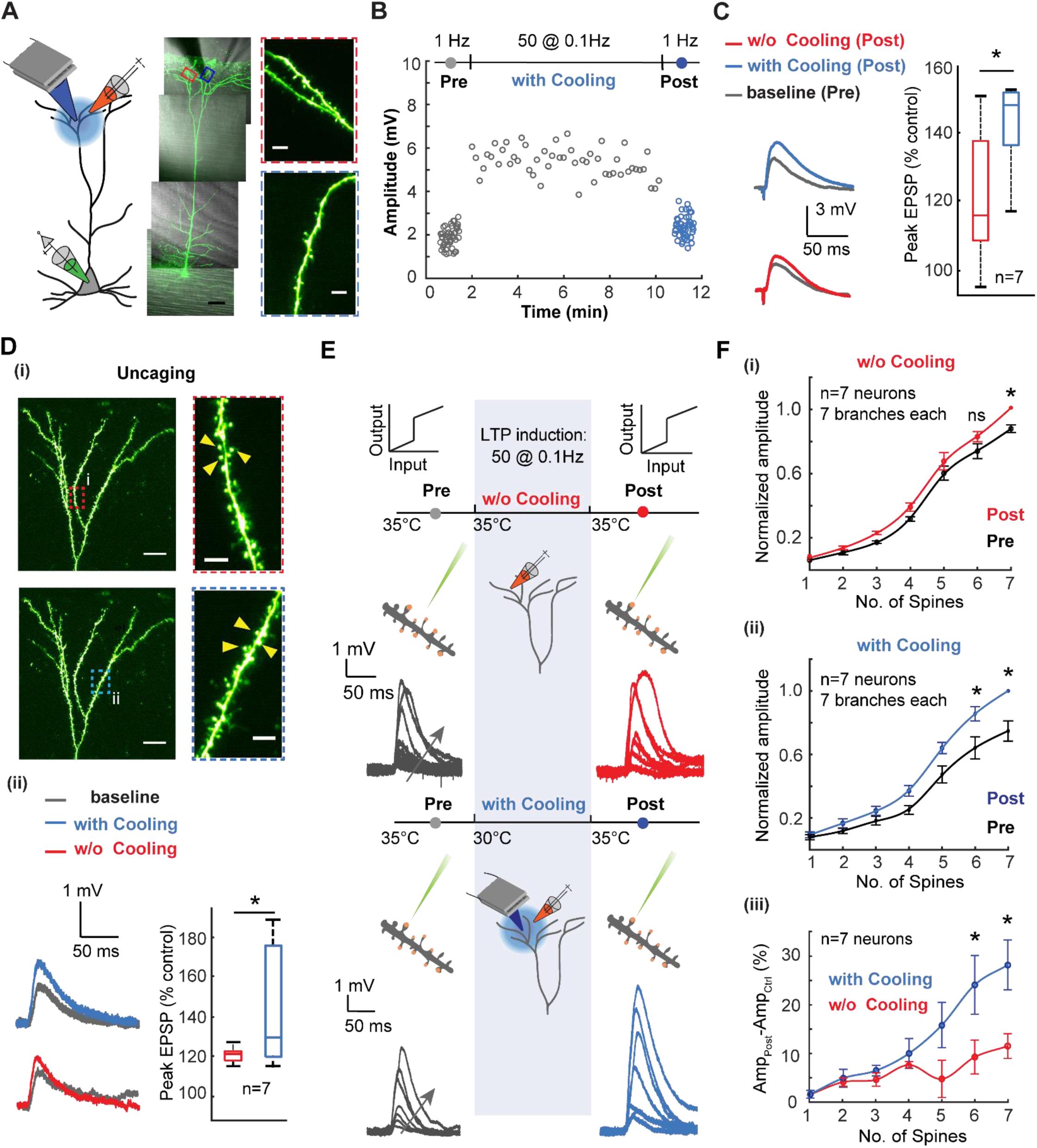
Mild focal cooling amplifies tuft dendritic plasticity and excitability. ***A)*** Schematic depicting low-frequency stimulation-induced plasticity in tuft dendrites under mild focal cooling (left). An exemplar layer 5 pyramidal neuron loaded with Alexa 488 (100 μM) (scale bars, 50 µm). (Inset) Dendritic segments that underwent the plasticity protocol without (top, red) and under the influence of focal cooling (bottom, blue) (scale bars, 5 µm). ***B)*** Field-evoked EPSP amplitude as a function of low-frequency plasticity induction in the presence and absence of mild focal temperature modulation (data not shown for the case without cooling). The initial readout at 1 Hz (gray) is followed by potentiation at 0.1 Hz and subsequent readout again at 1 Hz (blue). While the initial (grey) and final readouts (blue) are performed at 35^°^C, the plasticity induction is carried out at two different temperatures for comparison - 35^°^C (data not shown) and 30^°^C (grey). For clarity, the initial and final readouts at 1Hz and the potentiation (0.1 Hz) are only shown for the case in which plasticity was induced at 30^°^C. Note the induced plasticity is amplified under the influence of mild focal cooling. ***C)*** Comparison between the percentage of increase (post vs. control) in the field-evoked EPSP amplitude while low-frequency potentiation is performed at different temperatures (protocol as described in B). (Wilcoxon Mann Whitney test, n=7 neurons, p=0.0346). Error bars represent the standard error of the mean (SEM). ***D) (i)*** Uncaging-evoked responses from spines located on tuft dendritic branches plasticized using the plasticity protocol without the effect of cooling (top, red) and under the influence of cooling (bottom, blue) (scale bars, 20 µm). (Inset) Dendritic segments on which spines were uncaged before and after plasticity induction without (top, red) and under the influence of focal cooling (bottom, blue) (scale bars, 5 µm). **(ii)** Uncaging-evoked responses across spines located on target dendrites (left), depicting the difference between baseline (gray) and post-LTP induction (red: 35^°^C; blue: 30^°^C; signifies the temperatures at which plasticity was carried out), and percentage of increase (post-LTP vs. baseline control) for uncaging-evoked response (right) (Wilcoxon Mann Whitney test, n=7 neurons, p=0.0262). Error bars represent the standard error of the mean (SEM). ***E)*** Schematic of the experimental protocol for probing the temperature dependence of branch-specific potentiation via a readout of nonlinear input-output transformations across the tuft dendrite. Low-frequency field stimulation evokes plasticity, while two-photon glutamate uncaging probes nonlinear input-output characteristics before (control, grey) and after (post, either red or blue) plasticity induction. Arrows indicate an increasing number of spines stimulated. Focal cooling is only applied during the induction phase, while control and post-induction readout occur at 35^°^C. ***F)* (i)** Normalized dendritic input-output response measured at the soma before and after plasticity induction at 35^°^C (Wilcoxon signed rank test, n=7 neurons, *p<0.05). **(ii)** Similar to (i), but here plasticity induction was performed alongside moderate cooling (Wilcoxon signed rank test, n=7 neurons, *p<0.05). **(iii)** Comparison showing the percentage of amplitude increase for plasticity induced with focal cooling and without focal cooling (Wilcoxon Mann Whitney test, n=7 neurons, *p<0.05). Error bars represent the standard error of the mean (SEM).

To corroborate if the measured effect is indeed modifying the response post-synaptically, we performed pre- and post-induction readout using two-photon glutamate uncaging across spines (∼3 nearby spines; see Methods for details) located along several neighboring tuft dendritic branches at 35^°^C (**Fig. 1D (i)**). This readout was measured after a 10-minute plasticity induction protocol at either 35°C or 30°C. Dendrites that underwent the plasticity protocol under mild focal cooling exhibited a 44%±10% increase in the EPSP amplitude after low-frequency plasticity induction. In contrast, dendrites plasticized at the physiological baseline temperature reflected a 20%±1.4% increase in EPSP amplitude (Wilcoxon Mann Whitney test, n=7 neurons, p=0.0262) (**Fig. 1D (ii)**). Calcium dynamics triggered by glutamate uncaging also reflected an increase in dF/F (**see Methods**) after LTP is induced under mild cooling compared to the induction at physiological temperature (n=5 neurons, * p<0.05) (**Fig. S5**).

Next, we probed the temperature dependence of branch-specific excitation via a readout of nonlinear input-output transformations across the tuft dendrite (**Fig. 1E**). Here, low-frequency field stimulation evoked plasticity. At the same time, two-photon glutamate uncaging was used to probe nonlinear input-output characteristics before (control, grey) and after (post, either red or blue) plasticity induction, which was performed at either 35°C or under mild focal cooling at 30°C. Uncaging a sequence of 1 to 7 closely spaced dendritic spines (distributed along 10-20 µm stretches of a single branch) and recording the resulting EPSPs at the soma revealed a clear NMDA-dependent sigmoidal input-output transformation (**Fig. 1E,F**). This was followed by low-frequency plasticity induction with and without mild focal cooling and uncaging the same sequence again. It is critical to note that just as with field-evoked recordings, the pre-and post-induction readouts were performed at the physiological baseline of 35°C and the post-induction readout recorded at least ∼10 minutes after the low-frequency induction to ensure that the potentiation effect is not transient and a complete temperature recovery occurs (**see Fig. 1E**).

At physiological temperatures, i.e., without the application of mild focal cooling, tuft dendrite exhibited a moderate increase in the normalized EPSP amplitude after plasticity induction compared to their pre-induction baseline (∼ 11±2.6% under the condition in which seven spines were uncaged; Wilcoxon signed rank test, n=7 neurons, p=0.0156) (**Fig. 1F (i)**), while under moderate focal cooling, the increase in the EPSP amplitude was more pronounced with a 27.2±4.8% (under the condition in which seven spines were uncaged) increase compared to the pre-induction baseline (Wilcoxon signed rank test, n=7 neurons; p=0.0156). Thus, comparing the percentage of EPSP amplitude increase under conditions where plasticity induction was performed with and without cooling (**Fig. 1F (iii)**) reveals a significant effect of mild focal cooling on net dendritic plasticity and excitability (Wilcoxon Mann Whitney test, n=6 neurons, * p<0.05).

### Mild focal cooling does not impact basal dendritic plasticity and excitability

To compare the effect of thermal modulation between basal and tuft dendrites, we repeated the low-frequency plasticity induction experiments for basal dendrites (**Fig. 2A**). We once again performed field-evoked stimulation at 1 Hz as a baseline readout followed by a sequence of 50 pulses at 0.1 Hz then performed a post-induction readout at 1 Hz again. Similar to the experiments across the tuft dendrites, the initial (grey) and final readouts (either red or blue) were performed at 35^°^C. At the same time, the plasticity induction at 0.1 Hz was carried out at two different temperatures for comparison - 35^°^C (physiological baseline, data now shown) and 30^°^C (**Fig. 2B**). Care was taken to probe dendritic segments away from the soma (180-220 µm far from the soma) to avoid directly impacting spike output. We immediately noticed, and in stark contrast to the tuft dendrites, low-frequency field stimulation did not induce any amplification in the EPSP amplitude modulated during plasticity induction as a function of focal cooling (Wilcoxon Mann Whitney test, n=7 neurons, p=0.4136) (**Fig. 2C**). This effect was also mirrored in the uncaging-evoked responses across spines (**Fig. 2D (i)**), which did not reflect a significant change in the EPSP amplitude after low-frequency potentiation at both 35°C and 30°C (**Fig. 2D (ii)**) (uncaging-evoked, Wilcoxon Mann Whitney test, n=6 neurons, p=0.6282).

**Fig. 2.**
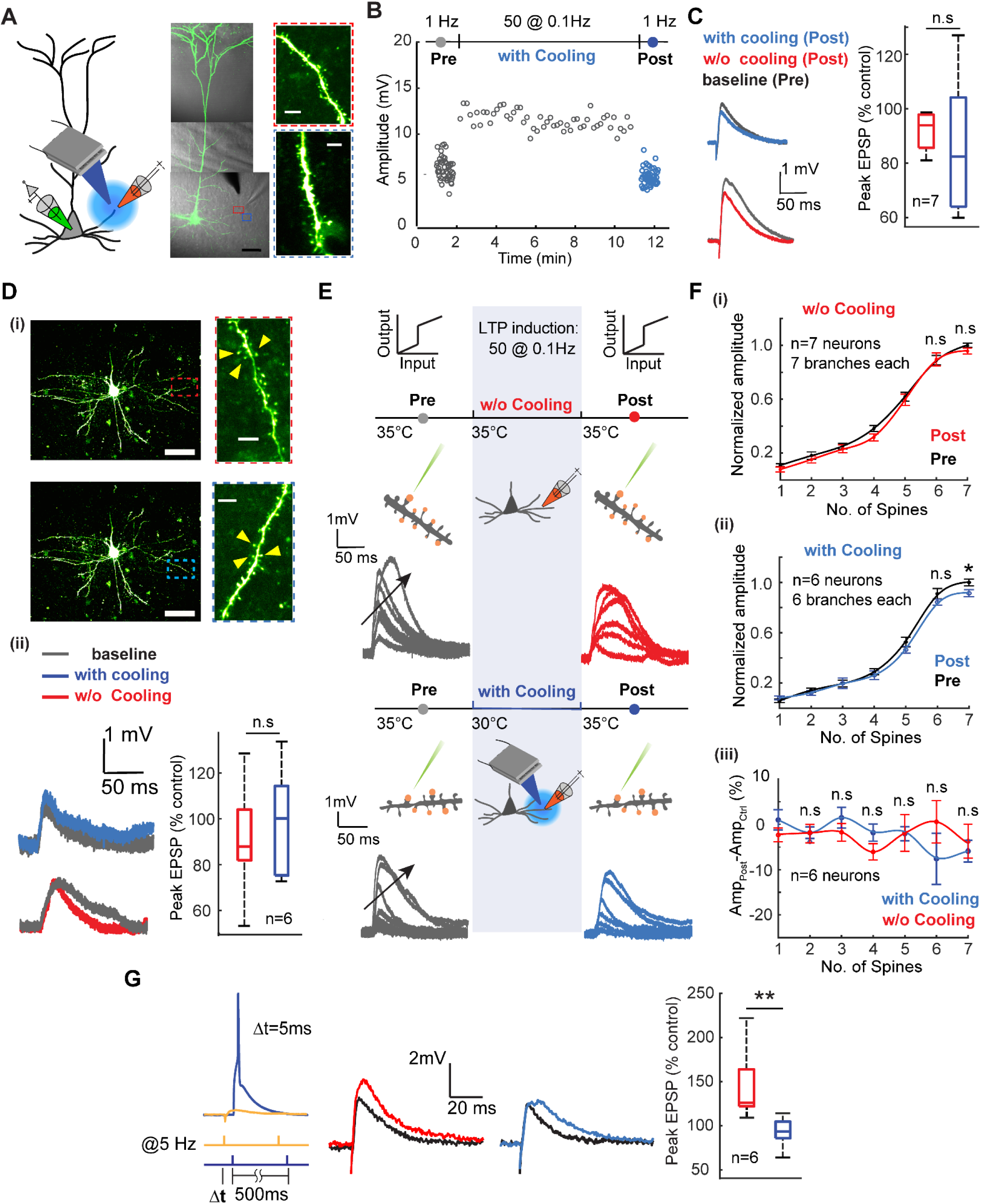
Mild focal cooling does not impact basal dendritic plasticity and excitability. ***A)*** Schematic depicting low-frequency stimulation-induced plasticity in basal dendrites under mild focal cooling (left). An exemplar layer 5 pyramidal neuron loaded with Alexa 488 (100 μM) (scale bars, 50 µm). (Inset) dendritic segments that underwent the plasticity protocol without (top, red) and under the influence of focal cooling (bottom, blue) (scale bars, 5 µm). ***B)*** Field-evoked EPSP amplitude as a function of low-frequency plasticity induction in the presence and absence of mild focal temperature modulation (data not shown for the case without cooling). The initial readout at 1 Hz (gray) is followed by potentiation at 0.1 Hz and subsequent readout again at 1 Hz (blue). While the initial (grey) and final readouts (blue) are performed at 35^°^C, the plasticity induction is carried out at two different temperatures for comparison - 35^°^C (data not shown) and 30^°^C (grey). For clarity, the initial and final readouts at 1Hz, as well as the potentiation (0.1 Hz) are only shown for the case in which plasticity was induced at 30^°^C. Note the induced plasticity is not significantly changed under the influence of mild focal cooling. ***C)*** Comparison between the percentage of increase (post vs. control) in the field-evoked EPSP amplitude while low-frequency potentiation is performed at different temperatures. Results of field stimulation show an insignificant change in the EPSP amplitude after LTP (Wilcoxon Mann Whitney test, n=7 neurons, p=0.4136). Error bars represent the standard error of the mean (SEM). ***D) (i)*** Uncaging-evoked responses from spines located on basal dendritic branches plasticized in the absence of cooling (top, red) and under the influence of cooling (bottom, blue) (scale bars, 50 µm). (Inset) Dendritic segments where the spines are uncaged before and after plasticity induction without (top, red) and under the influence of focal cooling (bottom, blue) (scale bars, 5 µm). **(ii)** Uncaging-evoked responses across spines located on target dendrites (left), depicting the difference between baseline (gray) and post-LTP induction (red: 35^°^C; blue: 30^°^C; signifies the temperatures at which plasticity was carried out), and percentage of increase (post-LTP vs. baseline control) for uncaging-evoked response (right) (Wilcoxon Mann Whitney test, n=6 neurons, p=0.6282). Error bars represent the standard error of the mean (SEM). ***E)*** Schematic of the experimental protocol for probing the temperature dependence of branch-specific potentiation via a readout of nonlinear input-output transformations across the basal dendrite. Low-frequency field stimulation is used to evoke plasticity, while two-photon glutamate uncaging probes nonlinear input-output characteristics before (control, grey) and after (post, either red or blue) plasticity induction. Arrows indicate increasing number of spines stimulated. Focal cooling is only applied during the induction phase, while control and post-induction readout occur at 35^°^C. Note the absence of amplification in the dendritic nonlinear response between the baseline (grey) and post-plasticity period (red, blue). Arrows indicate an increasing number of spines uncaged. ***F)* (i)** Normalized dendritic input-output response measured at the soma before and after plasticity induction at 35^°^C (Wilcoxon signed rank test, n=7 neurons, seven spines: p=0.3750, six spines: p=0.98125, five spines: p=0.3750, four spines: p=0.0313, three spines: p=0.2969, two spines: p=0.2969, one spine: p=0.1094). **(ii)** Similar to (i), but here plasticity induction was performed alongside moderate cooling (Wilcoxon signed rank test, n=6 neurons, seven spines: p=0.0303, six spines: p=0.4375, five spines: p=0.0313, four spines p=0.3125, three spines p=1.0, two spines p=0.3125, one spine p=0.3125). **(iii)** Comparison showing the percentage of amplitude increase for plasticity induced with focal cooling and without focal cooling (Wilcoxon Mann Whitney test, n=7 neurons, seven spines: p=0.5338, six spines: p=0.3660, five spines: p=0.7308, four spines: p=0.0734, three spines: p=0.5338, two spines: p=1.0, one spine: p=0.1014). Error bars represent the standard error of the mean (SEM). ***G)*** (left) As an additional control, a spike timing-dependent plasticity (STDP) protocol was used instead of low-frequency stimulation to evoke potentiation across basal dendrites with and without the application of mild focal cooling. (middle) Baseline (black) and post-induction readout without cooling (red) or with cooling (blue) applied. (right) Comparison of evoked amplitudes as a percentage of control for the two cases (Mann Whitney test, n=6 neurons, p = 0.002). A mild reduction in temperature during plasticity induction does not amplify excitability.

Basal dendrites are the locus of feedforward input in the cortex and are known to exhibit a myriad of nonlinear regenerative events that are both NMDA and Ca^2+^-dependent^53^. To examine the effect of focal cooling on the nonlinear integration in basal dendrites, we performed the local field-evoked stimulation at the dendrite with the previously established low-frequency plasticity protocol (a sequence of 50 EPSP at 0.1Hz) and readout using uncaging-evoked EPSPs (**Fig. 2E**). We first uncaged a sequence of synapses (distributed along 10-20 µm of a single branch). We recorded the resulting EPSPs at the soma as a function of an increasing the number of spines stimulated. We observed distinct nonlinear input-output transformations as a function of several synapses uncaged simultaneously (**Fig. 2E, F**), giving us a baseline input-output transformation. We induced plasticity via low-frequency stimulation at 35°C and 30°C, performed separately across different cells, and performed a post-induction readout at 35°C on the target branch. We observe that temperature did not modulate the input-output transformation either in slope, threshold, or amplitude in comparison to the effect observed in the tuft dendrite (**Fig.1**). The input-output curves for the condition in which no temperature modulation was applied during potentiation showed no change in the input-output curve (**Fig.2F (i)**). At the same time, the application of moderate focal cooling during the induction process resulted in a slight net decrease in the amplitude of the uncaging evoked EPSPs, especially once the number of spines needed for the nonlinear response passed the threshold (**Fig. 2F (ii)**) (n=6 neurons; one branch per neuron, p=0.0303). A comparison between potentiation with and without cooling shows mild focal cooling does not change the basal dendrite’s excitability (**Fig. 2F (iii)**). As an alternate form of plasticity and excitability, and to account for the frequency dependence of plasticity protocol in basal dendrites versus tuft dendrites, we used spike timing dependent plasticity (STDP) protocol^54,55^ to evoke potentiation across basal dendrites under cooling and without cooling (**Fig. 2G**). Here, the EPSP amplitude increased by 44% (144±17% of control) after STDP induction at 35°C, whereas the potentiation induced under mild focal cooling at 30°C did not exhibit any amplification but rather a slight depression (87±7.2% of control) in the EPSP amplitude (Wilcoxon Mann Whitney test, n=6 neurons, p=0.002).

### Mechanisms for cooling-modulated synaptic plasticity

As previously described, the Kv4.2 potassium channel is critical for the low-frequency induced plasticity observed in tuft dendrites. We hypothesized that the distribution of Kv4.2 potassium channels along the tuft dendrite and their dynamics as a function of mild cooling could be critical to the contrasting modulation observed between the apical tuft and basal dendrites. Given the lower temperature sensitivity of sodium channels^13^ compared to the higher temperature sensitivity of A-type potassium channels^22^, particularly Kv4.2, and the involvement of Kv4.2 channels in low-frequency-induced long-term potentiation (LTP)^48^, other types of LTP^56^, and branch-specific excitability in tuft dendrites^57^, our hypothesis is well grounded.

We applied 400 nM heteropodatoxin-2 to the bath solution to block Kv4.2 channels to test this hypothesis. Twenty minutes after application, we subjected the dendrites to a low-frequency stimulus protocol at two locations along the tuft dendrite—tertiary and primary branches. We recorded and averaged the field-evoked EPSPs before and after the low-frequency stimulation.

Our results revealed that tertiary dendrites were significantly affected by Kv4.2 channel blockade. Specifically, the change in EPSP amplitude before and after 0.1 Hz stimulation was 142% ± 13%, but this was reduced to 97% ± 10% after heteropodatoxin-2 application (Wilcoxon Mann Whitney test, n=5 neurons, p=0.046; **Fig. 3A**). In line with previous reports, this indicates that Kv4.2 channels are crucial for low-frequency-induced plasticity in tertiary tuft dendrites. In contrast, primary tuft dendrites did not show significant sensitivity to Kv4.2 channel blockade, with EPSP amplitudes changing from 123% ± 9% to 144% ± 21% after heteropodatoxin-2 application (Wilcoxon Mann Whitney test, n=5 neurons, p=0.9307; **Fig. 3B**).

**Fig. 3.**
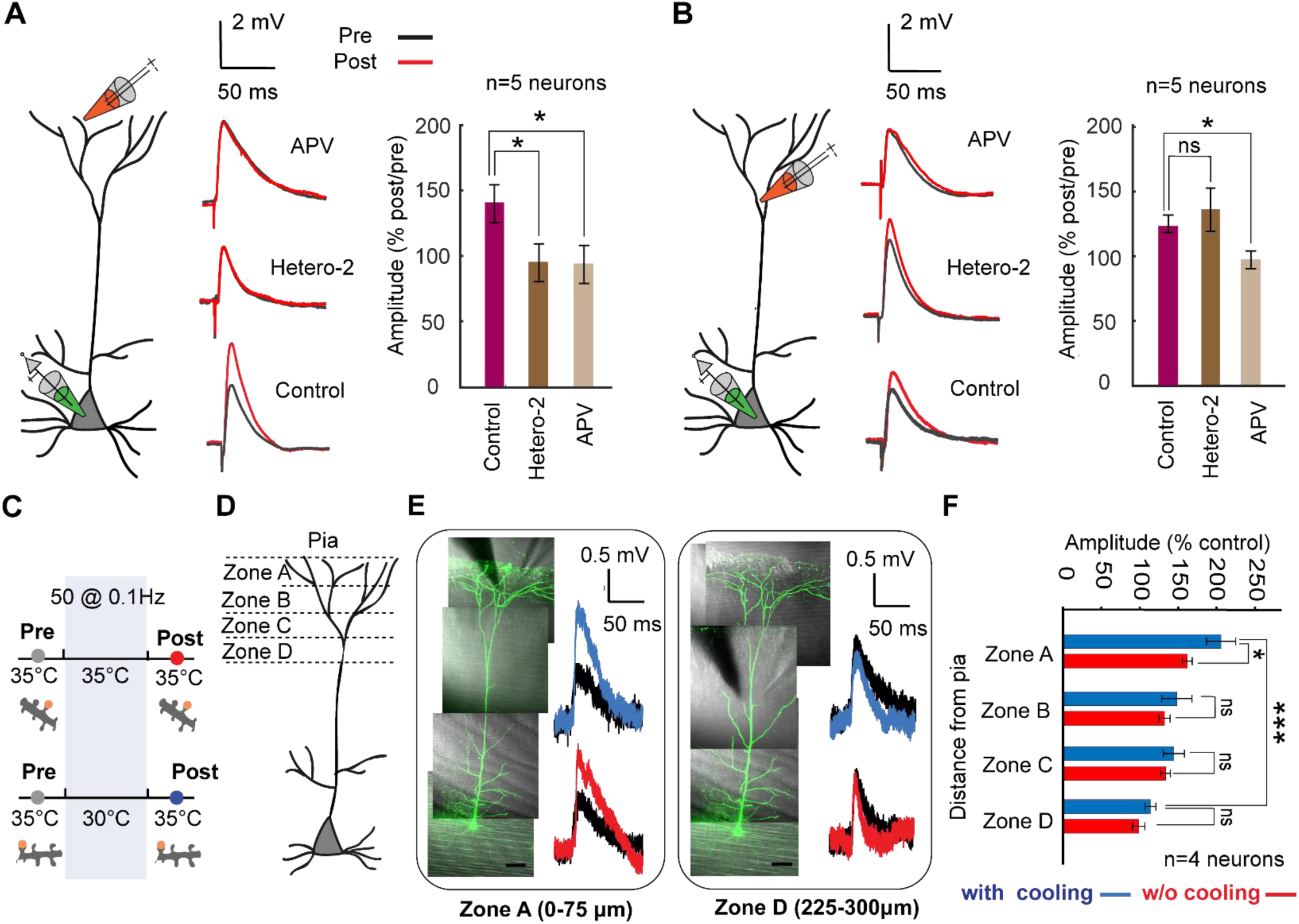
Gradient of plasticity along the apical dendrite. ***A)*** Experimental configuration. Field stimulation is used to evoke plasticity in the distal tuft dendrite (within 50 μm from pia) under the low frequency protocol. Example field-evoked EPSPs for pre (black) and post (red) stimulation for the control condition (bottom), after adding Heteropodatoxin-2 (Hetero-2, middle), and after adding APV (top). (Right) Comparison showing the percentage of EPSP increase (post vs. pre) with and without application of pharmacological blockers (Wilcoxon Mann Whitney test, n=5 neurons in each condition, Hetero-2 vs. control p=0.046 and APV vs. control: p=0.0159). Error bars represent the standard error of the mean (SEM). ***B)*** Similar experiment to that shown in (A) but performed in zone C at a different location of the tuft dendrite (within 200 μm from pia) (Wilcoxon Mann Whitney test, n= 5 neurons, Heter-2 vs. control: p= 0.9307, and APV vs. control: p=0.0159). Note the difference in both potassium channel dependence and amplification in plasticity in comparison to the distal tuft in (A). Error bars represent the standard error of the mean (SEM). ***C)*** Schematic of the experimental protocol for probing the gradient of the temperature dependence of low frequency induced plasticity along the tuft dendrite. Low-frequency field stimulation evokes plasticity, while two-photon glutamate uncaging is used to record the EPSP amplitude of control (grey) and after (post, either red or blue) plasticity induction. Focal cooling is only applied during the induction phase, while control and post-induction readout occur at 35^°^C. ***D)*** Schematic depicting different distal tuft dendritic zones (zone A: 0-75 µm from pia, zone B: 75-150 µm from pia, zone C: 150-225 µm from pia, zone D:225-300 µm from pia). ***E)*** Exemplar layer 5 pyramidal neurons (scale bars, 50 µm) shown for two different dendritic zones (zone A and zone D) that underwent field-evoked low-frequency plasticity. (Right) uncaging-evoked EPSPs depicting the difference between baseline (black) and post-LTP induction (red: 35^°^C; blue: 30^°^C; signifies the temperatures at which plasticity was carried out). ***F)*** Summary plot of the percentage of change in EPSP amplitude after LTP induction across different locations along the apical dendrite with and without mild focal cooling (Wilcoxon Mann Whitney test, n=6 dendrites from 4 neurons, *p<0.05, ***p<0.001). Error bars represent the standard error of the mean (SEM).

We also blocked N-methyl-D-aspartate receptors (NMDARs) in both dendritic branches by adding APV to the bath solution. The change in EPSP amplitude before and after potentiation was 96.1% ± 10% for tertiary dendrites (n=5, p=0.0159) and 92% ± 5.3 % for primary dendrites (n=5, p=0.0159) (**Figs. 3A, B**). These results suggest that NMDARs mediate low-frequency-induced potentiation in primary tuft dendrites, while NMDARs and Kv4.2 channels mediate potentiation in tertiary dendrites.

Our findings indicate that tertiary tuft dendrites likely have a higher density of Kv4.2 channels^58,59^ and exhibit greater temperature dependence in their plasticity responses than primary tuft dendrites.

The variability in cooling-enhanced excitability and low-frequency evoked plasticity along the tuft dendrite, combined with the uneven distribution of Kv4.2 channels along the apical dendrite^58^, led us to hypothesize a gradient of excitability and plasticity from the proximal to the distal end of the tuft. To investigate this, we induced low-frequency plasticity at two different temperatures and used glutamate uncaging to readout across single synapses located at varying distances from the pia: zone A (0-75 µm), zone B (75-150 µm), zone C (150-225 µm), and zone D (225-300 µm) (**Fig. 3C,D**).

Our results revealed a notable temperature dependence in EPSP amplitude changes in zone A. Specifically, in plasticity induction at 30°C, the change in EPSP amplitude due to potentiation was 203% ± 18.8%, significantly higher than the 159% ± 6.5% observed at 35°C (p=0.0401) (**Fig. 3E, F**). In contrast, no significant temperature-dependent differences were observed in zones B, C, and D. These findings highlight that low-frequency-induced plasticity is both temperature- and location-dependent along the tuft dendrite. For instance, at 30°C, the change in EPSP amplitude before and after potentiation was 203% ± 18.8% in zone A versus 111% ± 7.2% in zone D, demonstrating a significant spatial gradient (p=0.0007). This gradient substantiates the nonuniform distribution of Kv4.2 channels along the dendrite. Still, it sheds light on the impact of cooling activity from the brain surface, with the most distal dendritic regions of Layer 5 neurons being impacted the most.

Lower temperatures generally decrease the fluidity of the neuronal membrane^16,60^. This can affect the function and mobility of membrane proteins, including ion channels and receptors, potentially leading to altered synaptic transmission and neuronal excitability, including changes in capacitance^61,62^. Electrical capacitance is affected by a rapid rise in temperature, but evidence that mild gradual cooling modulates membrane capacitance similarly does not exist. Using voltage-clamp experiments to extract membrane capacitance confirmed that moderate cooling (to 30°C) did not alter or impact the membrane compared to 35°C (**Fig. S6**).

We also assayed how focal cooling impacts Ca²⁺ diffusion in the tuft and basal dendrites. By stimulating spines and shafts with a strong depolarizing pulse at 35°C, we observed Ca²⁺ transients in the shaft. Repeating this under moderate focal cooling of the tuft dendrite across the same segment revealed increased Ca²⁺ transients along short dendritic segments (∼10 µm), with greater non-uniformity in Ca²⁺ distribution under mild cooling (**Fig. S7**). This effect was absent in the basal dendrites. This suggests that cooling may reduce the ionic diffusion coefficient, leading to more localized Ca²⁺ accumulation. Such ionic diffusion effects could further enhance the nonlinear integration of synaptic inputs in the tuft, further exacerbating cross-talk and excitability.

### Somato-dendritic coincidence detection is impacted by tuft dendritic focal cooling

The non-uniform distribution of ion channels, such as Kv4.2, along the apical-tuft axis, not only creates gradients in excitability and plasticity but can also impact back-propagating action potentials (bAPs), thereby influencing the overall processing of neural information. As described in the sections above, mild focal cooling can accentuate these gradients, leading to more pronounced effects on how information is encoded and integrated along different parts of the dendrite. Within this framework, it is well known that Kv4.2 channels play a critical role in modulating distal dendritic excitability and backpropagating action potential frequency.

The observed cooling-induced depolarization and enhanced excitability of tuft dendrites, as described in Figure 1, led us to investigate whether selectively cooling these dendrites could improve the efficiency of bAPs along the somato-dendritic axis. This investigation is crucial for understanding neural coding, as it may modulate how feedback and feedforward inputs are processed across different cortical layers.

To explore this, we conducted whole-cell somato-dendritic patch-clamp recordings to map coupling between the soma and apical dendrite (200-300 µm from the soma) under the influence of mild focal cooling (**Fig. 4A**). We induced bAPs via somatic current injection and recorded the amplitudes both at the soma and dendrite under tuft cooling (at 30°C) and compared the responsivity to the same performed under physiological baseline (35°C). We noticed changes in both the amplitude and the integral area under the curve as a function of cooling (**Fig. 4B**). Specifically, we observed a significant increase in the first bAP spike amplitude from 35.5±3.86 mV at 35°C to 43±3.85 mV under cooling, reflecting a 25% average increase (Fig. 4B; n=8 neurons, p=0.0078). The integral area under the curve increased from 1.1 mV·s at 35°C to 1.3 mV·s at 30°C (n=8 neurons, p=0.0078).

**Fig. 4.**
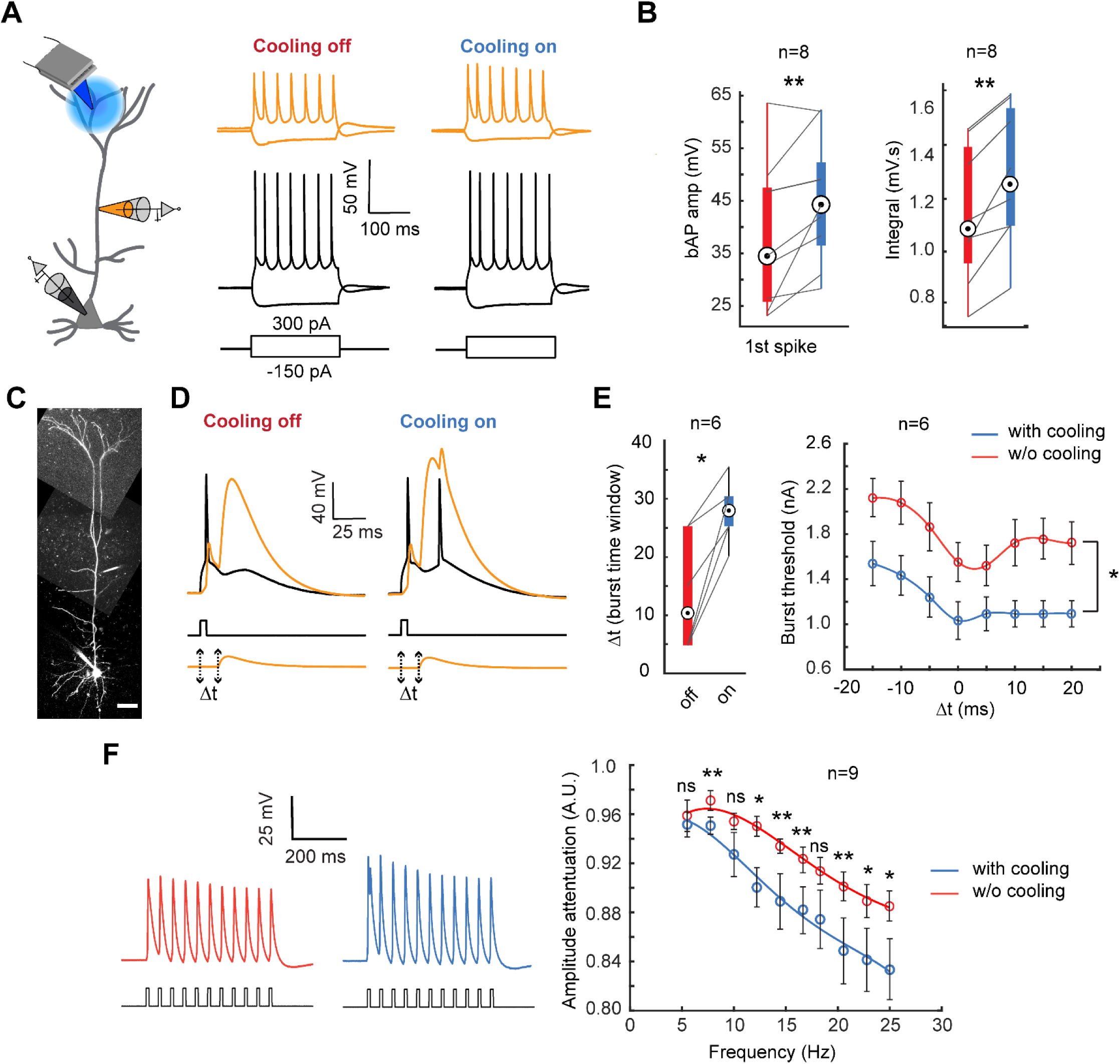
Somato-dendritic coupling modulation as a function of temperature. ***A)*** Schematic of the dual somatic and apical dendritic patch clamp recording while cooling the tuft dendrite (left). Prototypical spike trains recorded at the soma (bottom) and apical dendrite (top) under moderate focal cooling of the tuft dendrite at T=30°C (right) and T=35°C (left). ***B)*** The amplitude comparison of the first bAP spike (left) (Wilcoxon signed rank test, n=8 neurons, p=0.0078), and the integral (right) (Wilcoxon signed rank test, n=8 neurons, p=0.0039) comparing conditions with and without mild focal cooling. ***C)*** Exemplar layer 5 pyramidal neuron loaded with Alexa 488 (100 μM) under somato-dendritic patch clamping (scale bars, 50 µm). ***D)*** The combination of current injection at the soma and the apical dendrites EPSP-like depolarization separated by a time interval (Δt) of 10 ms evoked a burst following the onset of Ca^2+^ -AP in the apical dendrite while the tuft dendrite is focally cooled. ***E)*** The time duration (Δt) (left) and threshold needed to generate the burst (right) significantly increased and decreased, respectively, under mild focal cooling. (Wilcoxon signed rank test, n=6 neurons, *p<0.05). Error bars represent the standard error of the mean (SEM). ***F)*** The amplitude attenuation at the dendrite (ratio of the amplitude of the last spike to the third spike in a train of bAPs generated by 0.8 nA pulses at the soma) with and without mild focal cooling (Wilcoxon signed rank test, n=9 neurons for all points except for 10 Hz and 12.2 Hz where n=7, *p<0.05, **p<0.01). Error bars represent the standard error of the mean (SEM).

We next assayed if such a change leads to shifts in coincidence detection between bottom-up and top-down coupling evidenced via back-propagating action potential activated Ca^2+^ spike firing (BAC firing) (**Fig.4C, D**). Following the classical result of Larkum and colleagues^63^, combining a subthreshold EPSP-shaped distal dendritic potential and a back-propagating action potential within a time duration Δt elicited a Ca^2+^ and Na^+^ action potential complex in the dendrite. We observed an apparent increase in BAC firing efficiency as a function of cooling only the tuft dendrite (**Fig. 4D**). The time window (Δt) for burst induction and the threshold to generate the burst was more prolonged (13.5±4.01 ms without cooling compared to 27.5±2.14 ms under cooling; n=6 neurons; p=0.0313) and lower (Wilcoxon signed rank test, n=6 neurons, *p<0.05) under mild focal cooling of the tuft respectively (**Fig. 4E**). However, bAP amplitude attenuated more severely as a function of spike frequency under moderate focal cooling (**Fig. 4F**). We assayed this attenuation effect as follows. We measured as the ratio of the bAP amplitude of the last spike (in a train of eleven spikes) to that of the third spike. Our results indicate that bAP attenuation was more significant at 30°C, which became more pronounced with higher bAP frequencies (**Fig. 4F, right**). For instance, at 25 Hz, the spike amplitude showed 5% more attenuation (p=0.0391) compared to only 0.5% at 5.5 Hz (p=0.2754). This increased attenuation under cooling is most likely due to prolonged recovery from the inactivation of voltage-gated sodium channels as a function of temperature, which reduces bAP spike amplitude^64,65^. Together with the results from Figure 3, this would suggest a paradox wherein mild focal cooling, while depolarizing the tuft and increasing excitability and plasticity, reduces the efficacy of frequency-dependent bAP invasion, thereby controlling bursting. This differential effect and tradeoff are attributed to the differences in temperature-dependent dynamics of voltage-gated sodium and potassium channels.

### Neuronal coding is impacted by tuft dendritic focal cooling

Next, we examined how mild focal cooling of the tuft dendrite affects neuronal rate and burst coding, critical coding schemes^66–68^ in layer 5 pyramidal neurons, under an in-vivo-like conductance state. We used a custom-built dynamic clamp (**Fig. 5A, B**) (see **Methods**) and probed both somatic and dendritic outputs. We incorporated BAC firing into the dynamic clamp to ensure the output reflected bursts as would be observed *in vivo*. Bursts were identified when the inter-spike intervals (ISI) were shorter than 50 ms (**Fig. 5C**). Here we observe that while the average firing rate was not entirely perturbed, the occurrence of bursts reduced. As shown in **Fig. 5C**, a normalized ISI smaller than 0.15 indicated a burst. Under cooling, the burstiness (number of bursts divided by total spikes) averaged 0.17±0.04, significantly lower than the 0.26±0.07 observed without tuft cooling (**Fig. 5D, left**; p=0.0313). However, the mean ISI did not differ considerably between cooling and non-cooling conditions (**Fig. 5D, right**, p=0.6875). These results indicate that mild focal cooling of the tuft reduces burstiness but does not significantly affect the average ISI. This suggests a mechanism through which top-down integration could be selectively modulated in isolation.

**Fig. 5.**
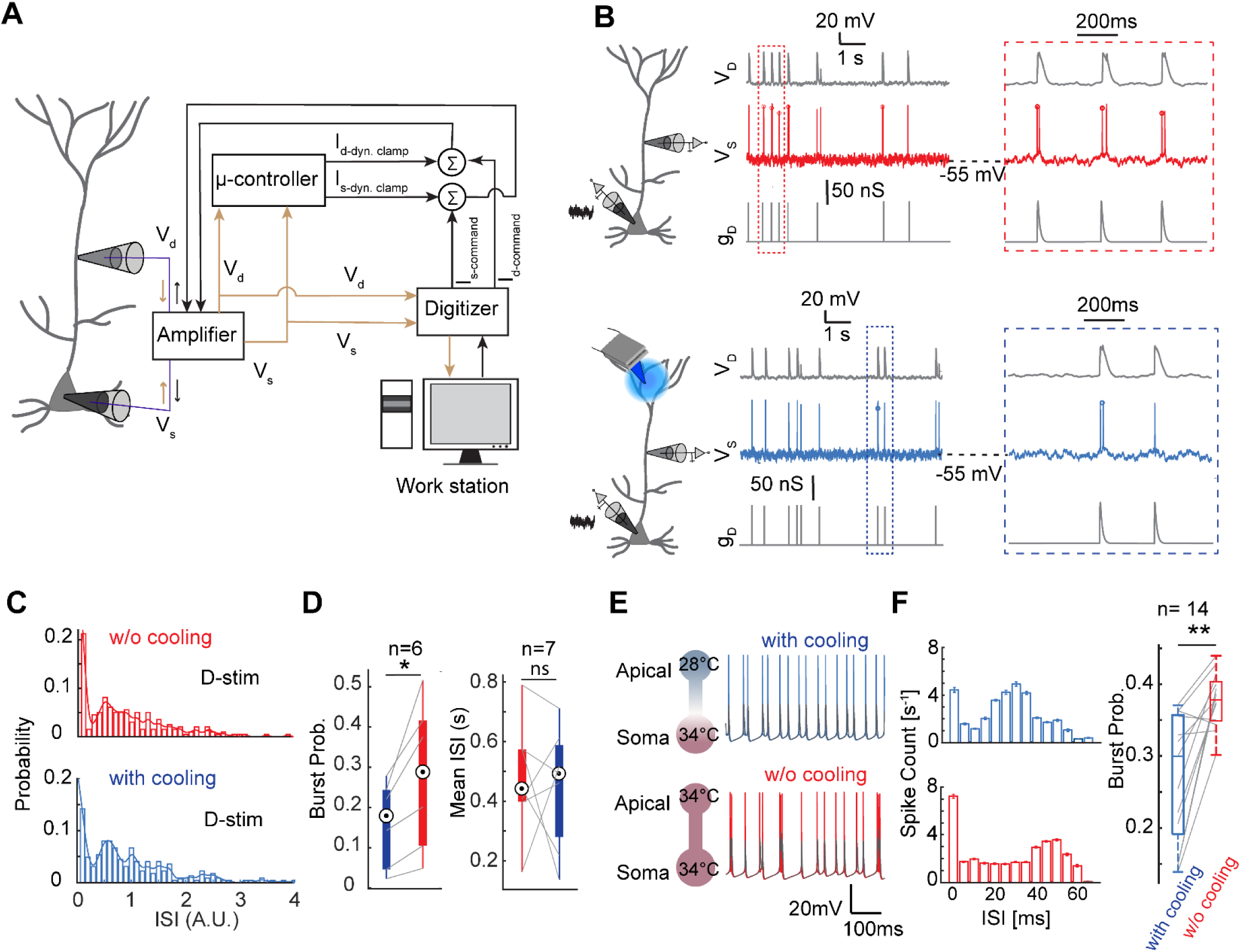
Temperature modulation across the tuft dendrite causes somatic rate regulation. ***A)*** Schematic of the dynamic clamp circuit that generates in-vivo-like conductance state through a feedback loop. ***B)*** Cooling effect on the intrinsic burstiness when a subset of bAPs is paired with apical excitatory conductance without (top block) and with cooling (bottom block). ***C)*** Histogram of the probability of the normalized I.S.I of the spikes in the presence of dendritic input without and with cooling. ***D)*** The ratio of the number of the burst to the total number of the spikes defined as burst probability is presented as function of the temperature when a dendritic input is coincident with a bAP (Wilcoxon signed rank test, n=6 neurons, p=0.0313) and the mean ISI (Wilcoxon signed rank test, n=7 neurons, p=0.6875). Note: a burst is defined as two consecutive spikes with the ISI smaller than 50 ms. ***E)*** Spiking activities of a three-compartment biophysical model under apical dendritic focal cooling (top) and physiological temperature (bottom), gray traces indicate corresponding dendritic voltage. ***F)*** Burst probability is reduced under apical focal cooling compared to physiological temperature (n = 14, the variance is due to imposed modulation in the dendritic geometry across models, see Methods for details, Wilcoxon signed rank test, p < 0.005).

To mechanistically corroborate this effect, we resorted to computational modeling (**Fig. 5E-F**) using a multicompartment model of a layer 5 thick-tufted neuron^69^ (see **Methods, and Fig. S8A**). In the model, we incorporated long-term inactivation kinetics of apical dendritic Na^+^ channels and their temperature sensitivities^64^, and recapitulated the temperature dependence of frequency-dependent attenuation of bAP amplitude (**Fig. S8A-C**). To validate the temperature sensitivity in this model, we cooled the perisomatic compartment and observed a reduction in spike rate as described in previous literature^12,13,52^ (**Fig. S8D**). The three-compartment biophysical model shows that focal cooling of the apical tuft dendrites alters the output spiking pattern compared to physiological temperature conditions (**Fig. 5E**). We observed that the mean firing rate did not appear to be significantly different (cooling: 34.23 ± 0.8635 Hz, physiological baseline temperature: 33.1077 ± 1.0283 Hz, mean ± std), but the burstiness (**Fig. 5F**) was impacted considerably even with a mere 5° to 6°C difference between tuft and soma (n = 14 (the variance is due to imposed modulations in the dendritic geometry across models, see Methods for details), Wilcoxon signed rank test, p<0.005). Critically, this effect was significantly impacted by the gating kinetics of Na^+^ channels and the temperature dependence of recovery from inactivation (**Fig. S8B-D**). When the temperature dependence of only Na^+^ channels was removed, the reduction in burstiness was abolished and increased above the baseline (**Fig. S8E**). Together, this establishes a model in which K^+^ channels in the tuft are responsible for the increased excitability. Still, the Na^+^ channel’s dependence on temperature via their recovery from inactivation controls the burst code and output rate. This indicates that a temperature-induced degeneracy^70,71^ between K^+^ and Na^+^ controls L5 pyramidal output, resulting in an apparent reduction in output gain and the appearance of slowing cortical output.

## DISCUSSION

In this study, we explore the nuanced effects of mild focal cooling on the dendritic properties of layer 5 intrinsically bursting pyramidal neurons (L5PNs), focusing on how focal cooling impacts the excitability and plasticity of these neurons at a single-cell level. Layer 5 pyramidal neurons are exciting because they serve as the principal output layer of the cortex^72^ and feature distinct dendritic compartments: apical tuft and basal dendrites, which receive top-down and bottom-up inputs, respectively^46,63,73–75^. Our study shows how mild focal cooling influences dendritic plasticity and excitability in L5PNs, with distinct effects observed in tuft versus basal dendrites. The results advance our understanding of dendritic plasticity and hold potential implications for neural modulation and therapeutic interventions. This is critical given that mild focal cooling applied at the cortical surface is generally used to study timing and slow down neural activity. Our results are some of the first steps in understanding how cell types in the cortex respond to such focal applications. As described, we find that the nonhomogeneous temperature profile across L5PNs confers differential tuning of dendritic action.

First, we demonstrated that mild focal cooling (from 35°C to 30°C) amplifies dendritic excitability and low-frequency induced plasticity, specifically in tuft dendrites of L5 pyramidal neurons. This amplification is evident from the significant increase in EPSP amplitude during and after low-frequency stimulation (**Fig. 1**). At baseline temperature (35°C), the amplitude increase was 22%, while it was 49% under cooling. This suggests that cooling enhances the synaptic efficacy and long-term potentiation (LTP) in tuft dendrites, a finding supported by two-photon glutamate uncaging experiments. Notably, the persistence of increased EPSP amplitude long after the cooling has been removed (observed at 35°C) indicates that the effects of cooling on plasticity are not transient (**Fig. 1** and Supplementary **Fig. S4**). This highlights the potential for thermal modulation to induce lasting changes in synaptic strength. Probing nonlinear input-output transformations revealed that cooling enhances this characteristic in tuft dendrites. The increased EPSP amplitude under cooling conditions demonstrated a more pronounced NMDA-dependent sigmoidal transformation, indicating that the cooling effect is likely due to enhanced NMDA receptor activity or associated signaling pathways, a feature critical for local plasticity and excitability.

In stark contrast to tuft dendrites, basal dendrites did not exhibit any significant change in plasticity or excitability under mild focal cooling and low-frequency induction (**Fig. 2**). We further confirmed the lack of cooling effect on basal dendrites using a spike-timing-dependent plasticity (STDP) protocol^55^, where no significant improvement was observed. These findings underscore the distinct functional roles of tuft and basal dendrites. Basal dendrites, which handle feedforward input and exhibit complex nonlinear integrative properties^11,41^, seem less sensitive to temperature modulation regarding plasticity enhancement and, in general, are less adaptable to temperature changes than those in tuft dendrites. Overall, the above results postulate that feedforward sensory drive might be less impacted by temperature modulation, but top-down feedback signaling contextual integration, which usually occurs in the tuft region, might be more impacted.

The tuft and basal dendritic compartments have different neuronal integration and plasticity roles. Apical tuft dendrites, which are somewhat isolated from the soma and other dendritic compartments, are characterized by a higher density of voltage-gated potassium channels than basal dendrites^58,76,77^. This unique property results in less excitability in apical dendrites and attenuates bAPs as they travel toward the tuft bifurcation. The diminished bAPs lead to reduced calcium electrogenesis, which is crucial for long-term potentiation (LTP)^45,78^, and can instead promote long-term depression (LTD)^32,46^. However, when the apical dendrites are depolarized during neuronal firing, it is possible to generate regenerative calcium activity that can switch plasticity from LTD to LTP^46^. Our observations using mild focal cooling of the tuft suggest that such a condition could be engineered via subtle cooling, leading to a new way of neuromodulation, even potentially creating a switch from LTD to LTP. Especially given that we observe an increase in bAP amplitude and increased excitability to unitary inputs, it would suggest increased soma-apical coupling – a feature we show via both amplitude and threshold of activation of BAC firing. That said, the same cannot be said about the reactivation of bursts within a window.

Probing the channel mechanisms, we find that Kv4.2 channels, known for their high sensitivity to temperature changes^22^, are crucial for low-frequency-induced plasticity in tuft dendrites. We demonstrate that blocking these channels with heteropodatoxin-2 impairs plasticity, specifically in distal tuft dendrites (**Fig. 3**). At the same time, more proximal apical regions remain unaffected, highlighting a spatial gradient in Kv4.2 channel density and functionality along the tuft-apical axis. This differential sensitivity suggests that distal tuft dendrites rely more on Kv4.2 channels for plasticity and might be the underlying differential response across the basal dendrites observed.

Next, we employed dual somato-dendritic patch clamping and targeted focal cooling of the tuft to investigate differences in coincidence detection and neural coding (**Fig. 4**). We found that moderate temperature drops of approximately 5°C enhance the amplitude of bAPs and calcium plateau potentials in apical tuft dendrites, indicating increased top-down excitability. Notably, we observed that cooling increases the likelihood of burst generation when the timing between somatic and dendritic inputs is less synchronized. From a coding perspective, this implies that cooling the superficial layer may enhance the integration and coincidence detection of delayed inputs, potentially improving input-output transformation.

Paradoxically, however, while the calcium plateau and bAP amplitude appear to be enhanced by cooling, the rate of recovery from inactivation for sodium channels seems to be impaired (**Fig. 5 and Fig. S8**), evidenced by a decay in bAP amplitude as a function of frequency, leading to a reduction in axo-somatic output^65,79^. This reveals a tradeoff between Kv4.2 and Na^+^ channel responsivity across the tuft-apical axis as a function of local temperature impacting neural coding and information integration. Such regulation leading to a reduction in burst coding might manifest across several downstream brain regions and appear to slow down circuit function. Our study suggests that while the overall somatic output code might reflect a reduction in excitability, local properties in the tuft signal an excitation increase. The current understanding is that a reduction in neuronal excitability can impair the ability of neurons to participate in plasticity processes. For example, diminished excitability may hinder the formation of new synaptic connections or the modification of existing ones, affecting learning and memory^80^. Our recordings suggest that the tuft is more excitable and resistant to this effect under mild focal cooling, but the same cannot be said about the basal dendrites. Overall, our study provides a unique method and mechanism through which one could differentially control top-down vs bottom-up integration in terms of timing with analog control.

## MATERIALS AND METHODS

### Acute slice preparation

All experimental procedures were conducted in accordance with the guidelines set forth by the NIH and Purdue Institutional Animal Care and Use Committee (IACUC). All physiological solutions were fully oxygenated (95% O_2_ and 5% CO_2_), pH adjusted to 7.3 to 7.4, and osmolarity maintained between 300 to 310 mOsm unless stated. Adult C57BL/6 (Jackson Laboratory) mice (both male and female, 8 to 12 weeks of age) were deeply anesthetized with 3 to 4% isoflurane followed by trans-cardiac perfusion with ice-cold NMDG cutting solution^81^ consisting of (in mM): 92 NMDG, 30 NaHCO_3_, 1.2 NaH_2_PO_4_, 20 HEPES, 2.5 KCl, 25 glucose, 5 sodium ascorbate, 3 sodium pyruvate, 2 thiourea, 0.5 CaCl_2_, 10 MgCl_2_, 5 N-acetyl-L-cysteine, before decapitation. Coronal slices (300-350 µm thickness) were prepared using a vibratome (Leica VT1200S) in 0° to 4° Celsius NMDG cutting solution. Brain slices were then allowed to recover in 34°C NMDG cutting solution with gradual perfusion of 0.25 to 0.5 ml Na^+^ rich NMDG solution (2M NaCl in NMDG cutting solution) over 6 to 10 minutes depending on the mouse age^82^ and transferred to room temperature HEPES artificial cerebrospinal fluid (ACSF) holding solution consisting of (in mM): 92 NaCl, 2.5 KCl, 1.25 NaH_2_PO_4_, 30 NaHCO_3_, 20 HEPES, 25 glucose, 5 sodium ascorbate, 3 sodium pyruvate, 2 thiourea, 2 CaCl_2_, 2 MgCl_2_, 5 N-acetylL-cysteine for at least 1h prior to recording.

### Electrophysiological recording

Slice recordings followed protocols previously established^41^. In short, slices were transferred to a chamber, continuously superfused with oxygenated ACSF, and visualized with an upright two-photon microscope (Bruker Nano, Madison, WI) comprising of an Olympus BX51WI body (Olympus, Tokyo, Japan) fitted with infra-red (IR) Dodt-gradient-contrast (DGC) optics, an IR sensitive camera (IR-2000, Dage-MTI, Michigan City, IN), and a 40× water immersion objective (0.8 NA, Nikon USA). Recordings were performed in 35°C recording ACSF consisting of (in mM): 125 NaCl, 3 KCl, 25 NaHCO_3_, 1.25 NaH_2_PO_4_, 25 glucose, 3 sodium pyruvate, 1 sodium ascorbate, 1.3 CaCl_2_, 1 MgCl_2_. 4 to 8 MΩ borosilicate patch pipette (Sutter Instruments, CA, USA) was pulled using a P1000 pipette puller (Sutter Instruments, Novato, CA), filled with internal solution containing (in mM): 130 potassium gluconate, 7 KCl, 10 HEPES, 5 NaCl, 35 sucrose, 2 MgSO_4_, 2 sodium pyruvate, 4 Mg-ATP, 0.4 Tris GTP, 7 phosphocreatine disodium (pH 7.3, osmolarity 290mOsm). 25 µM Alexa 594 or 100 µM Alexa 488 was used for two-photon structural imaging, 200 µM fluo-4 was used for two-photon Ca^2+^ imaging. L5 neurons were patch-clamped and had resting membrane potentials between -60 mV and -70 mV at rest without any current injection. Currents of ∼-50 pA to -100pA was injected in cases where the RMP was more negative (-75 mV) or needed to be maintained. Whole-cell recordings were made using a Multiclamp 700B, (Molecular devices, San Jose, CA), Bessel-filtered at 4kHz, and digitized at 4 to 20 kHz using a Digidata 1550B interface (Molecular devices, San Jose, CA) and winwcp software.

### Focal cooling

The cooling device is consisted of a 250µm diameter silver wire (A-M systems, #781000; thermal conductivity 406 W/mK) wrapped with 25µm graphene sheet (graphene-supermarket: Conductive Graphene Sheets, thickness: 25 mm; thermal conductivity: 1300-1500W/m in x-y plane and 13-15W/m in z plane) and insulated with a polyimide tube (1.1 mm diameter, Accu-Glass Product Inc., #111217)^83^. The polyimide tube was further sealed with adhesive epoxy (Newark, TBS20S; thermal conductivity: 1.1 W/m.K) in both ends to create a contained air insulation around the silver wire and avoid ACSF leakage into the tube. Only 2 mm of the silver wire was protruding out of the insulation layers to be exposed for cooling the tissue. The silver wire was coiled on one end and glued with Arctic Silver Adhesive (Custom Thermoelectric, TG-AS5-12G) to the cold side of a Peltier device (Custom Thermoelectric, 02301-9B30-32RU6A) which was itself getting cooled on its hot side by a homemade aluminum active heat sink. To have a better spatial resolution in cooling the spots on the tissue, the tip of the silver probe was sharpened by electro etching with sodium nitride solution to deliver a tip size of 10-30 µm. Prior to starting electrophysiological experiments, a k-type temperature sensor (Omega, 80 mm wires, product number 5SC-TT-K-40-72) was used to calibrate the temperature of the probe tip and the tissue once the probe was into the tissue. The solution temperature in the chamber was stable at 34.9 °C and the tip could be cooled down to the 29.6 °C in the best case.

### Thermal Transport Simulation

Numerical simulations were performed to model a two-dimensional thermal transport for the probe passing through ASCF and inserted in the slice using a finite element based COMSOL Multiphysics 5.6. The effect of the diameter and length of the silver wire were investigated to find the optimal design for cooling the slice and the results are summarized in Fig. S2.

### Field Evoked Stimulation

Focal electrical field evoked stimulation was performed via a theta-glass (borosilicate; Hilgenberg) pipette pulled using a P1000 pipette puller (Sutter Instruments, Novato, CA), and filled with the recording ACSF solution. The pipette tip was located at approximately 5-10 µm from the target dendritic segment guided by the fluorescent image of the dendrite. A biphasic current pulse with 0.1 ms duration was injected through the electrode at various intensities (1–5 µA) via stimulus isolator (ISO-Flex; AMPI). The effectiveness and localization of the stimulation were verified by measuring the Ca^2+^ intensity resulting from the stimulus pulses.

### Pharmacology

For the plasticity experiments, gabazine (500µM; Sigma) and metabotropic gamma-aminobutyric acid receptor blockers (10 mM CGP-55845) were added to the recording ACSF solution. In specific experiments, NMDAR blocker (100 mM APV; Tocris Bioscience) and Kv4.2 subunit channel blocker (0.5 mM heteropodatoxin-2; Alomone) were added to the ACSF perfusion solution 20 minutes before the start of the recording session. After the experiments, the recording ACSF solution containing the pharmacological blockers was washed in through a perfusion system, and the next recording session was performed 10-20 minutes after the pharmacological blocker wash-in.

### Dynamic Clamp

A microcontroller-based circuit board was used for performing dynamic clamp experiments as previously described^41^. The module was controlled with custom-written codes (Arduino, Processing). First, the linear input-output relationship of the amplification circuit was measured with a DC voltage source (E3631A Agilent Technologies, Santa Clara, CA) and oscilloscope (Keysight, Santa Rosa, CA), and then verified with a model cell (Molecular devices, San Jose, CA). The working principle is as follows: the microcontroller reads the output of the amplifier, and then A/D converts the amplified membrane voltage, computes the current based on differential equation models, and D/A converts the current, which is then summed with the current command from the Digidata 1550B interface (Molecular devices, San Jose, CA). The ODEs are solved using the forward Euler method.

The Ornstein-Uhlenbeck process-based point conductance model^84^ mimics background synaptic inputs under a noisy in vivo-like state. We used this point conductance model instead of a Poisson train of synaptic inputs^85^ to ensure higher variability in the amplitude of EPSPs and IPSPs, better reflecting the background synaptic inputs with different synaptic strengths. The dynamic conductance is computed with the following differential equations (Equations 1, 2, and 3) (χ_1_ and χ_2_ are two independent random variables following unit normal distribution)

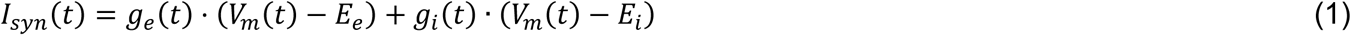

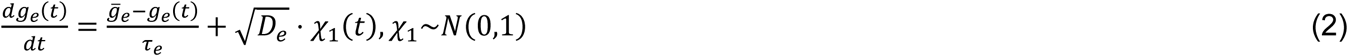

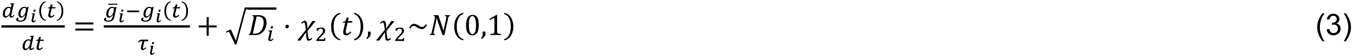

In our experiment, τ_*e*_ = 2.8 ms, τ_i_ = 8.5 ms, E_e_ = 0 mV, E_i_ = −80 mV, were chosen based on the physiological properties of α-amino-3-hydroxy-5-methyl-4-isoxazolepropionic acid (AMPA) and γ-Aminobutyric acid (GABA) channels. The mean of noisy excitatory conductance (g_e_) varies between 2 nS to 4 nS for different amount of spontaneous background firing rate, and g_i_ equaled 2 nS to 9 nS. Conductance values agreed with previous literature and this range helped match the background firing rates to that observed in vivo (2–20 Hz), but avoided a hyperexcitable state. The noise diffusion coefficients of the excitatory (D_e_) and inhibitory conductance (D_i_) were scaled to match voltage fluctuations normally observed in vivo (∼5 mV RMS). These terms, which contribute to the “noisy” nature of dynamic conductance, were in line with previous literature^84^ and this ensured the fast-fluctuations in the background were preserved.

To mimic coincident apical synaptic inputs and bAPs (Figure 5A&B), a double exponential apical excitatory conductance (G_max_ = 50 to 70 nS, E_e_ = 0 mV, τ_1_ = 1 ms, τ_2_ = 10 ms) was applied through the dendritic dynamic clamp once the dendritic voltage exceeded the bAP threshold (-40 mV to -15 mV dependent on the dendritic patch location) in a close-loop manner.

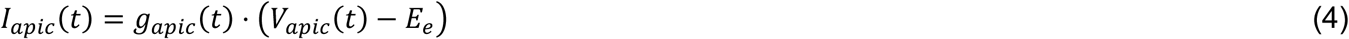

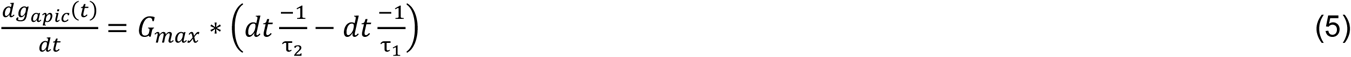

### Two-photon imaging and glutamate uncaging

A laser-scanning microscope (Bruker Nano, Madison, WI) with dual galvanometric mirrors and two femtosecond pulsed lasers (Spectra Physics, Insight X3 and Mai-Tai) were used to perform two-photon imaging and glutamate uncaging. Laser beam intensities were independently controlled with electro-optical modulators (model 350-50; Conoptics, USA). The imaging beam (set to 810 nm for Alexa 594 and fluo-4, and 920 nm for Alexa 488) was merged with the uncaging beam (720 nm) using a 760 nm long-pass dichroic mirror. During Calcium (Ca^2+^ imaging, line scans were performed across the spine head and the adjacent branch with 8–10 μs dwell time (400–800 Hz line rate) and 3 mW–5 mW laser power at the objective focal plane. Structural imaging was performed at 5–8 μs dwell time. For glutamate uncaging with 4-Methoxy-7-nitroindolinyl-caged-L-glutamate (MNNI-glu, Tocris, MN, USA), 2.5 mM MNI-glu was diluted in freshly prepared recording ACSF and applied to the bath through a circulating pump. At these concentrations, there was no epileptiform-like activity. The uncaging dwell time was 0.5 ms and the laser power needed for uncaging ranged from 20 mW to 30 mW. At these powers, no visible photodamage occurred. Baseline fluorescence of both channels was continuously measured to assay any damage. Ca^2+^ transients were also measured to ensure spines were still functionally active with no loss in physiological response. EPSP time course and changes in resting membrane potential following repeated stimulation were also assayed as indicators of any photodamage. Table-top hard shutters were used to avoid exposure and any off-target uncaging. Ca^2+^ signals were expressed as DF/F (calculated as (F - F_baseline_)/F_baseline_)^41^. Data was collected from dendrites at least 30 μm below the surface of the slice and were not prematurely cut off before termination.

### Three-compartment biophysical model

The simplified biophysical model of L5 PNs consisted of three different compartments: soma-AIS, proximal apical dendrite and distal apical dendrite. The ion channels and corresponding parameters in each compartment can be found in Table S1 for more details. To mimic the diversity of morphology and intrinsic ion channel distribution across the L5 PN population, the ratio between the total membrane surface area of each compartment, as well as the distance between the dendritic compartments to soma, were assigned with 5% jitter around the average value in **Table S1**. The distance between proximal apical dendrite and soma was kept larger than 210 µm to ensure intrinsic burstiness. Bursts were defined as two or more spikes occurring with 20 ms time interval or less. The differential equations were solved with the forward Euler method with an interval of 0.05 ms. The dendritic Na^+^ channel (nad) model was adapted from previous studies^64,65^, the Q10 value was set to 2.3 for all ion channels. The code implementing this simulation is available on github: https://github.com/shulanx1/simpl5pn_temperature_modulation.

## Data, Materials, and Software Availability

Upon acceptance, all data will be uploaded to Zenodo, and code will also be uploaded to GitHub repositories wherever applicable.

## ACKNOWLEDGMENTS

This work is based upon efforts supported by EMBRIO Institute, contract #2120200, a National Science Foundation (NSF) Biology Integration Institute to K.J. This work was also supported by the following grants to K.J. NIH R21EB029740 Trailblazer Award; Human Frontiers Science Program (HFSP) RGY0069; ORAU Ralph E. Powe Junior Faculty Enhancement Award, the National Institutes of Health, New Innovator Award NIH DP2MH136494. This material is also based upon work supported by the Air Force Office of Scientific Research under award numbers FA9550-22-1-0078 and FA9550-23-1-0701. Any opinions, findings, and conclusions or recommendations expressed in this material are those of the author(s) and do not necessarily reflect the views of the United States Air Force. M.H.M was supported by the NSF-ASEE E-Fellows program

## Author Contributions

M.H.M., S.X., and K.J. designed the study and experiments. M.H.M. performed all experiments. S.X. assisted with experiments. M.H.M., S.X., and K.J. analyzed and interpreted data. M.H.M. and K.J. wrote the manuscript. All authors edited the manuscript. K.J. provided overall guidance and funding for the study.

**Competing interest**: The authors declare no competing interest.

## Key Resource Table

| REAGENT or RESOURCE | SOURCE | IDENTIFIER |
| --- | --- | --- |
| Chemicals, peptides, and recombinant proteins |  |  |
| Gabazine | Sigma Aldrich | SR95531; CAS: 152622-30-0 |
| D-AP5 | Tocris | 0106; CAS: 79055-68-8 |
| Heteropodatoxin-2 | Alomone | H-154; CAS:152939-43-6 |
| Alexa Fluor 594 Hydrazide | Invitrogen | A10438 |
| Alexa Fluor 488 Hydrazide | Invitrogen | A10436 |
| Fluo-4, Pentapotassium Salt, cell impermeant | Invitrogen | F14200 |
| Sodium chloride | Sigma-Aldrich | S9888; CAS: 7647-14-5 |
| Potassium chloride | Sigma-Aldrich | P9333; CAS: 7447-40-7 |
| Sodium bicarbonate | Sigma-Aldrich | S5761; CAS: 144-55-8 |
| Sodium phosphate monobasic | Sigma-Aldrich | S8282; CAS: 7558-80-7 |
| HEPES | Sigma-Aldrich | H4034; CAS: 7365-45-9 |
| Glucose | Sigma-Aldrich | G8270; CAS: 50-99-7 |
| Sucrose | Sigma-Aldrich | S0389; CAS: 57-50-1 |
| Calcium chloride | Sigma-Aldrich | C8106; CAS: 10035-04-8 |
| Magnesium chloride hexahydrate | Sigma-Aldrich | M2670; CAS: 7791-18-6 |
| Adenosine 5'-triphosphate magnesium salt | Sigma-Aldrich | A9187; CAS: 74804-12-9 |
| Guanosine 5'-triphosphate tris salt | Sigma-Aldrich | G9002; CAS: 103192-46-7 |
| Phosphocreatine disodium salt hydrate | Sigma-Aldrich | P7936; CAS: 19333-65-4 |
| Potassium gluconate | Sigma-Aldrich | P1847; CAS: 299-27-4 |
| N-Acetyl-L-cysteine | Sigma-Aldrich | A7250; CAS: 616-91-1 |
| N-Methyl-D-glucamine | Sigma-Aldrich | 66930; CAS: 6284-40-8 |
| Sodium hydroxide | Sigma-Aldrich | S8045; CAS: 1310-73-2 |
| Potassium hydroxide | Sigma-Aldrich | 221473; CAS: 1310-58-3 |
| Sodium L-ascorbate | Sigma-Aldrich | A4034; CAS: 134-03-2 |
| Sodium pyruvate | Sigma-Aldrich | P5280; CAS: 113-24-6 |
| Thiourea | Sigma-Aldrich | T8656; CAS: 62-56-6 |
| Hydrochloric acid | Sigma-Aldrich | 320331; CAS: 7647-01-0 |
| Deposited data |  |  |
| Primary data | This paper | https://doi.org/10.5281/zenodo.14020289 |
| Experimental models: Organisms/strains |  |  |
| Mouse: C57BL/6J | The Jackson Laboratory | IMSR_JAX:000664 |
| Software and algorithms |  |  |
| Biophysical simulation | This paper | [DOI: 10.5281/zenodo.14020643](https://doi.org/10.5281/zenodo.14020643) |
| Prairie view | Bruker | NA |
| MATLAB (R2018a) | MathWorks | https://www.mathworks.com |
| COMSOL Multiphysics | COMSOL, Inc. | Version 5.6 |
| NEURON (7.6) | Hines and Carnevale<sup>86</sup> | https://neuron.yale.edu/neuron/ |
| ImageJ | Schneider et al.<sup>87</sup> | https://imagej.nih.gov/ij/index.html |
| Arduino | Arduino | https://www.arduino.cc/ |
| Materials and reagents |  |  |
| Silver wire | A-M systems | #781000 |
| Graphene sheet | graphene-supermarket | SKU: GRAPH-SHEET-25-8X4-5P |
| polyimide tube | Accu-Glass Product Inc | #111217 |
| adhesive epoxy | Newark | TBS20S |
| TG-AS5-12G | Custom Thermoelectric | TG-AS5-12G |
| Thermoelectric Cooler | 02301-9B30-32RU6A | 02301-9B30-32RU6A |
| k-type Temperature sensor | Omega | 5SC-TT-K-40-72 |
| Other |  |  |
| Dynamic clamp board | This paper | https://doi.org/10.5281/zenodo.11303939 |
| Ultima *In Vitro* multiphoton microscope | Bruker Nano | SCR_017142 |
| 512 × 512 Spatial Light Modulator | Meadowlark Optics | NA |
| MaiTai HP | Spectra-physics | NA |
| InSight X3 | Spectra-physics | NA |
| Prism compressor | Newport | PC-TS-KT |
| Objective | Nikon | N40X-NIR |
| Vibratome | Leica | VT1200 S |
| Multiclamp 700B | Molecular devices | NA |
| Digidata 1550B | Molecular devices | NA |
| ISO-Flex Stimulator | AMPI | https://www.ampi.co.il/iso-flex |

