## Supporting information for "Mild focal cooling selectively impacts computations in dendritic trees"

**This PDF file includes:**

Figures S1 to S8

Tables S1

SI References

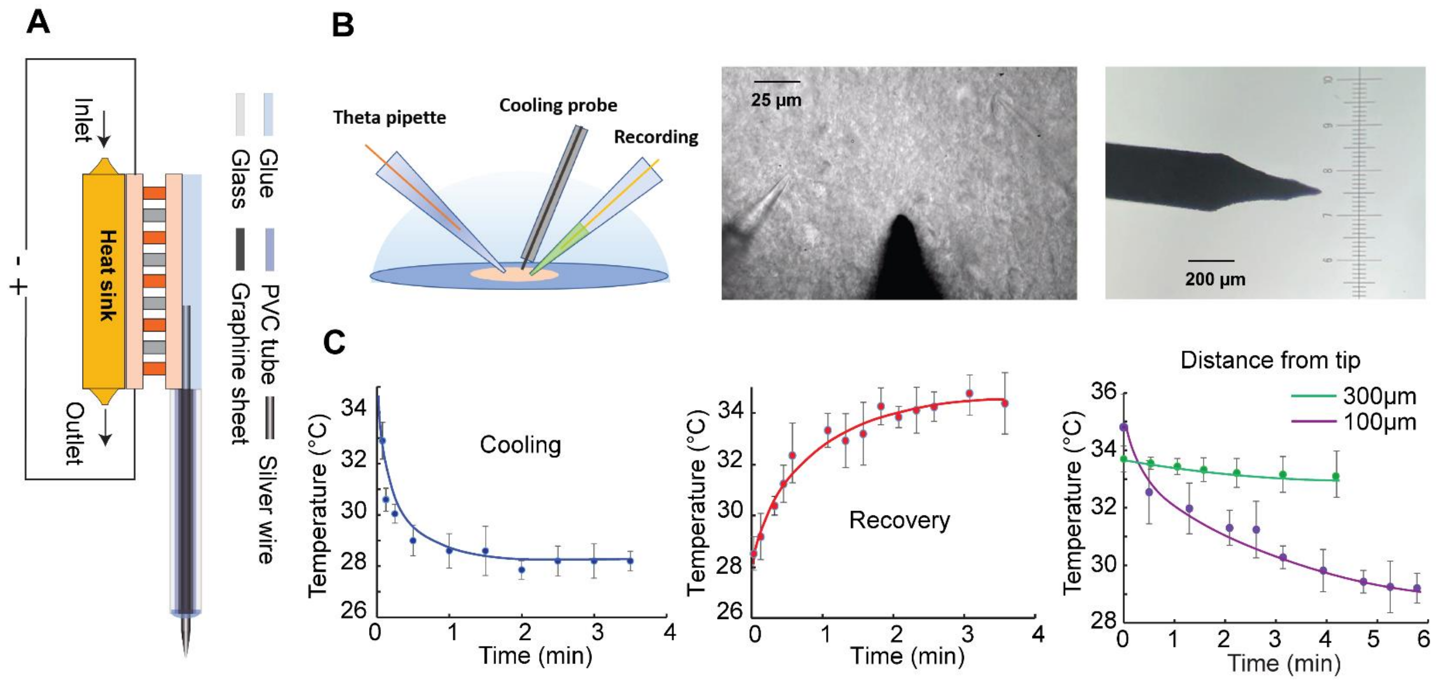


***Fig. S1*. *Custom-designed cooling probe and temperature calibration***

***A)*** *Schematic of the focal cooling device,* ***B)*** *Schematic of the recording and stimulation pipettes, and cooling probe on the slice (left), the Dodt contrast of the probe and pipettes on the brain slice (middle), and the probe tip under the microscope.* ***C)*** *Temperature measured with the thermocouple for cooling (left ), recovery (middle), and at different distances from the tip of cooling probe (right). It takes about 2 minutes for the temperature in the slice to reach the steady state condition, and 3 to 4 minutes for the cooled section of the slice to recover to its original temperature. Note: At a distance of ~300 μm from where the cooling probe tip is, the drop in cooling is not significant, reflecting a ~ 0.5°C drop in the temperature; however, at a 100 μm distance from the probe tip, the temperature drop is substantial (5-6°C).*

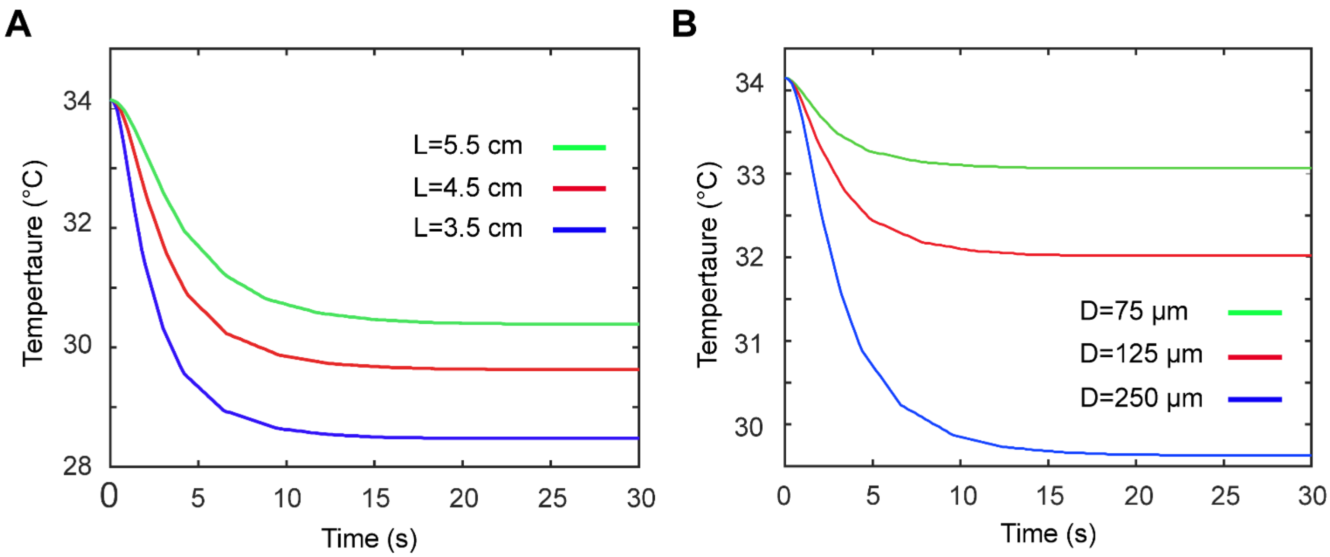


***Fig. S2. Simulation results for focal cooling of a spot on the slice in a recording chamber***

*The temperature at the tip of the silver probe as a function of time for various silver probe lengths* ***(A)*** *and different probe diameters* ***(B)****.*

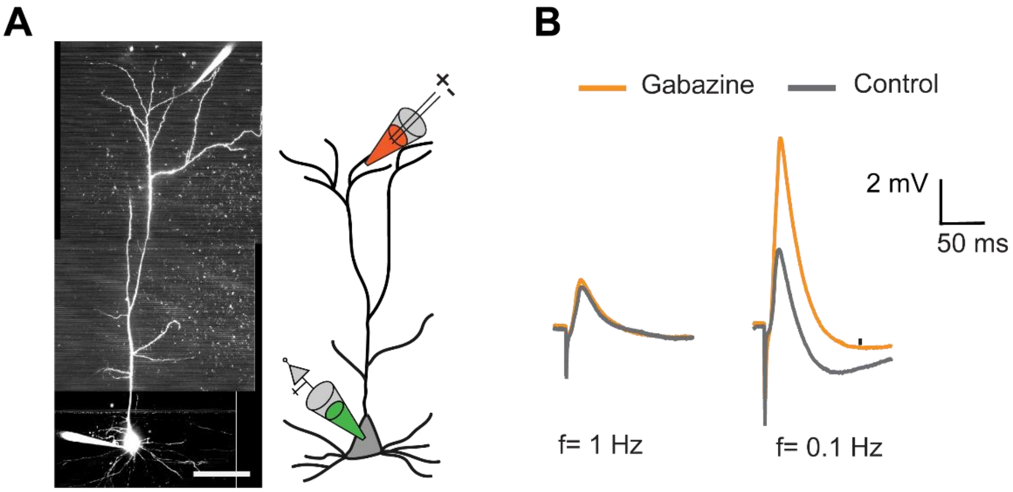


***Fig. S3. Gabazine blocks the inhibitions for low-frequency stimulation***

***A)*** *Fluorescence image of a layer 5 pyramidal neuron loaded with Alexa 488 (100 µM) and the pipettes' positions schematic.* ***B)*** *Example EPSPs for the control experiment and after adding Gabazine are presented for different frequencies of field stimulation.*

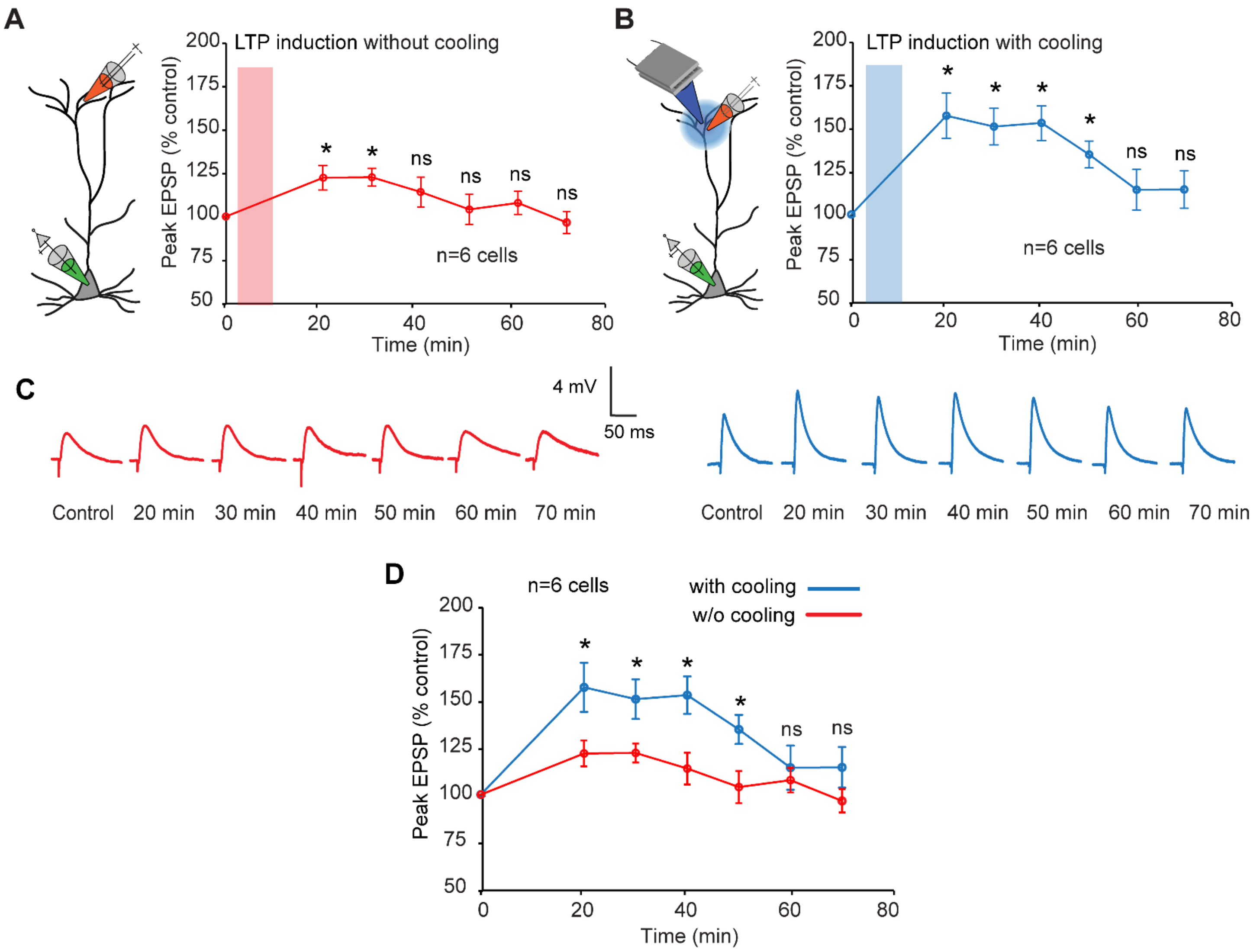


**Fig. S4. Time course of the low frequency induced synaptic plasticity at the tuft dendrite**

*Field-evoked EPSP amplitude as a function of low-frequency plasticity induction in the presence and absence of mild focal temperature modulation. The initial readout at 1 Hz is followed by potentiation at 0.1 Hz and subsequent readouts at 1 Hz and at different times after the low-frequency induction. While the initial (pre) and post-induction readouts are performed at 35°C, the plasticity induction is carried out either at 35°C (for the "without cooling" experiment) (A) or 30°C for the experiment with cooling (B).* ***A)*** *Percentage increase (post-LTP vs. baseline control) of field-evoked EPSP amplitude recorded every 10 minutes after the LTP induction at physiological temperature (Wilcoxon signed rank test, n=6 neurons, *p<0.05). (Error bars represent the standard error of the mean (SEM)).* ***B)*** *The same results as in A in the presence of cooling while LTP induction is performed.* ***C)*** *Example EPSP at different points of time for the plasticity induction in the absence (red) or presence of (blue) of cooling.* ***D)*** *Comparison between the time course of the plasticity effect for the induction at 35°C (red) and 30°C (blue) (Wilcoxon Mann Whitney test, n=6 neurons, *p<0.05). Error bars represent the standard error of the mean (SEM).*

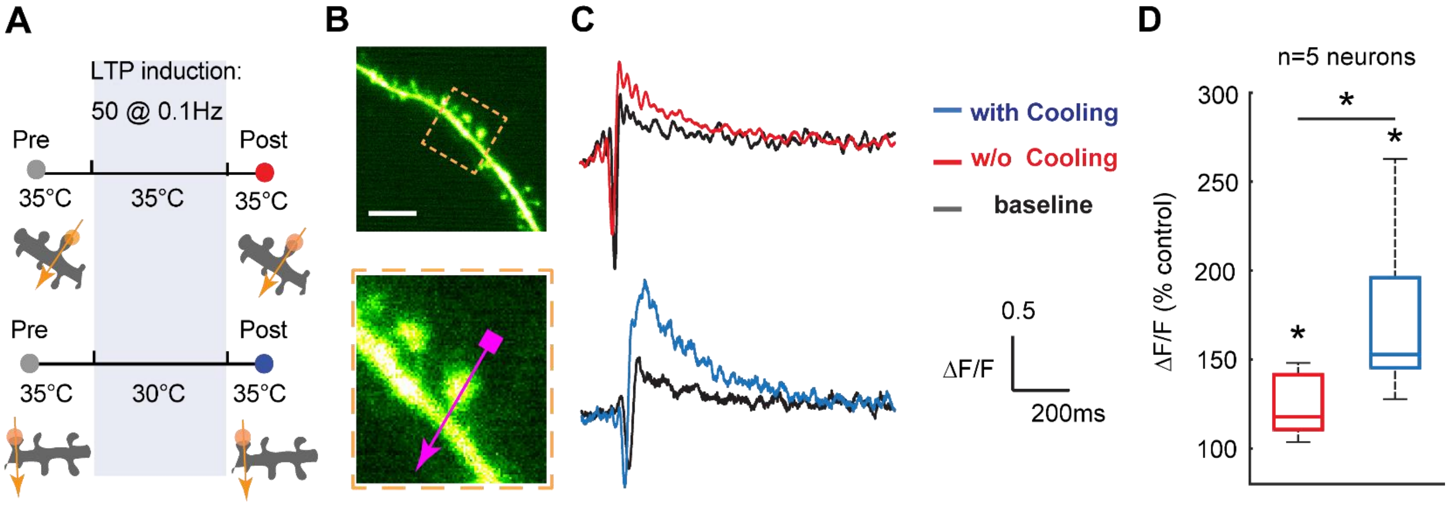


***Fig. S5. Mild focal cooling amplifies calcium transients in tuft dendrite***

***A)*** *Schematic of the experimental protocol for probing the temperature dependence of low frequency induced plasticity at the tuft dendrite. Low-frequency field stimulation is used to induce plasticity. Two-photon calcium imaging triggered with spine uncaging is used to record the calcium transients before (control, grey) and after (post, either red or blue) plasticity induction. Focal cooling is only applied during the induction phase, while control and post-induction readout occur at 35°C.* ***B)*** *Fluorescence image of the spine undergoing line scan calcium imaging (top) and the zoomed view (bottom). (Scale bar 5 μm).* ***C)*** *Exemplar calcium transients showing the baseline (black) and post-LTP induction (red: 35°C; blue: 30°C; signifies the temperatures at which plasticity was carried out).* ***D)*** *Percentage change (post-LTP vs. baseline control) of uncaging-evoked ΔF/F at the spine stimulated with mild focal cooling and without cooling* (Wilcoxon signed rank test, n=5 neurons*, *p<0.05*) and comparison between the two (Wilcoxon Mann Whitney test, n=5 neurons*, *p<0.05*)*. Error bars represent the standard error of the mean (SEM).*

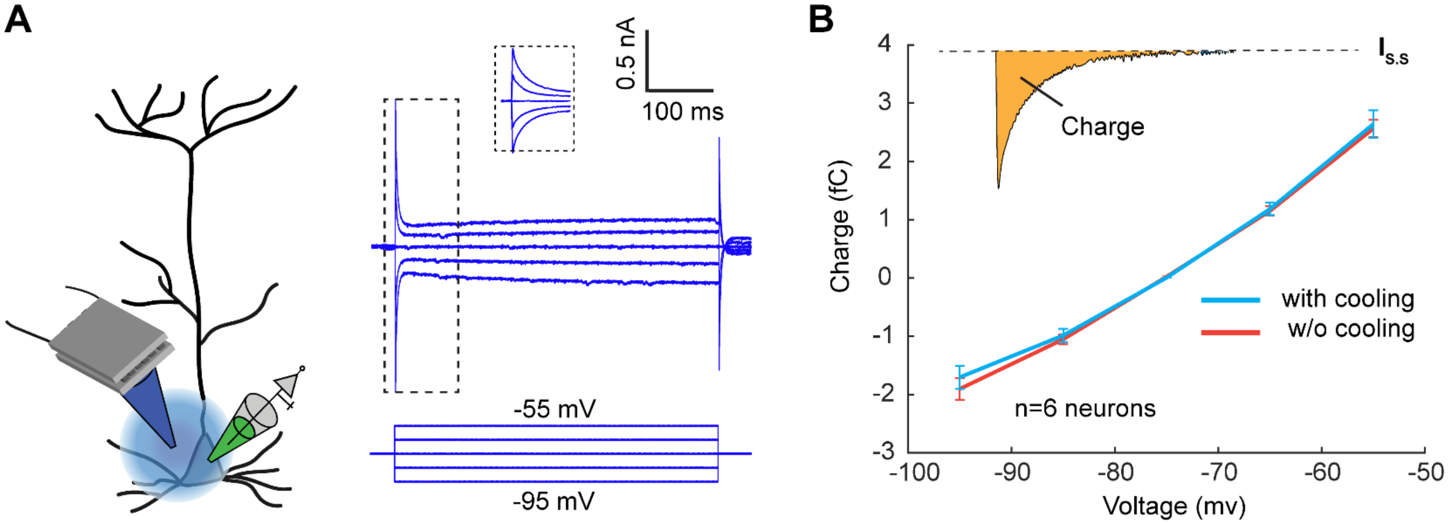


***Fig. S6. Electrical capacitance (charge) as a function of voltage for different temperatures***

***A)*** *Voltage clamp somatic recording is used to measure the charges through current-associated charge accumulation in the cell membrane. The voltage pulses are given in 5 steps from -95 mV to -55 mV (below the firing threshold) to allow measuring the capacity (or ion charges).* ***B)*** *Ionic charge as a function of the voltage with and without cooling**.** For the charge measurement, the equation* $Q = \int I(t)dt$ *is calculated from the time the pulse is initiated until the current reaches the steady state (*$I_{SS}$*), as shown in the inset. We do not observe any significant change in capacitance, possibly because the rapid temperature gradient alters electrical capacitance, not the temperature itself* [1,2]*.*

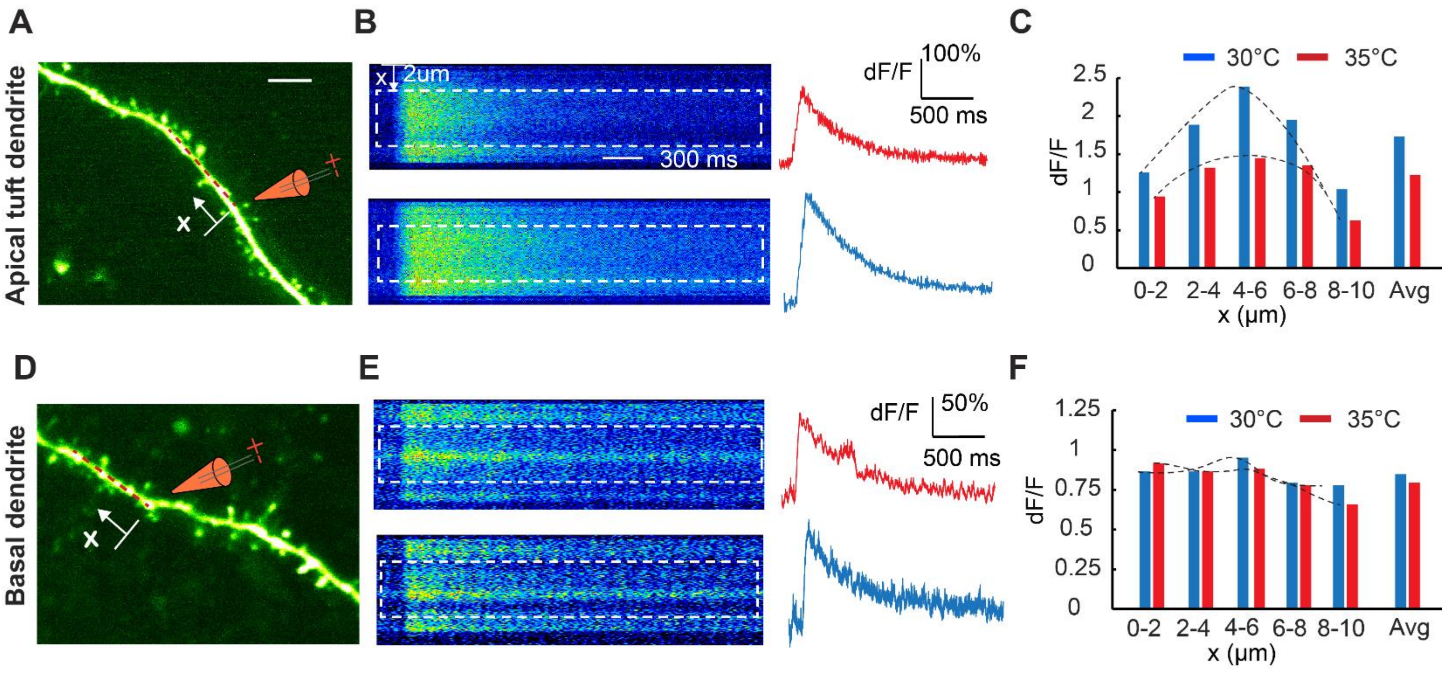


***Fig. S7. Temperature dependence of Ca$^{2+}$ diffusion along basal and apical dendrites***

***A)*** *Two-photon fluorescence image of the segment of apical tuft dendrite undergoing field-evoked stimulation. Scale bar: 5 µm.* ***B)*** *Ca transient along 10 µm segment of the tuft dendrite resulted from the single pulse field-evoked stimulation at 35°C (top) and 30°C (bottom), and the average calcium transient (dF/F) of the dashed segment of the line scans for the corresponding segments with and without cooling (red: 35°C, blue: 30°C).* ***C)*** *Calcium transients as a function of x (length of the segment) showing a higher amplitude under cooling than near physiological temperature.* ***D-F)*** *A similar experiment was performed on the basal dendrite segment.*

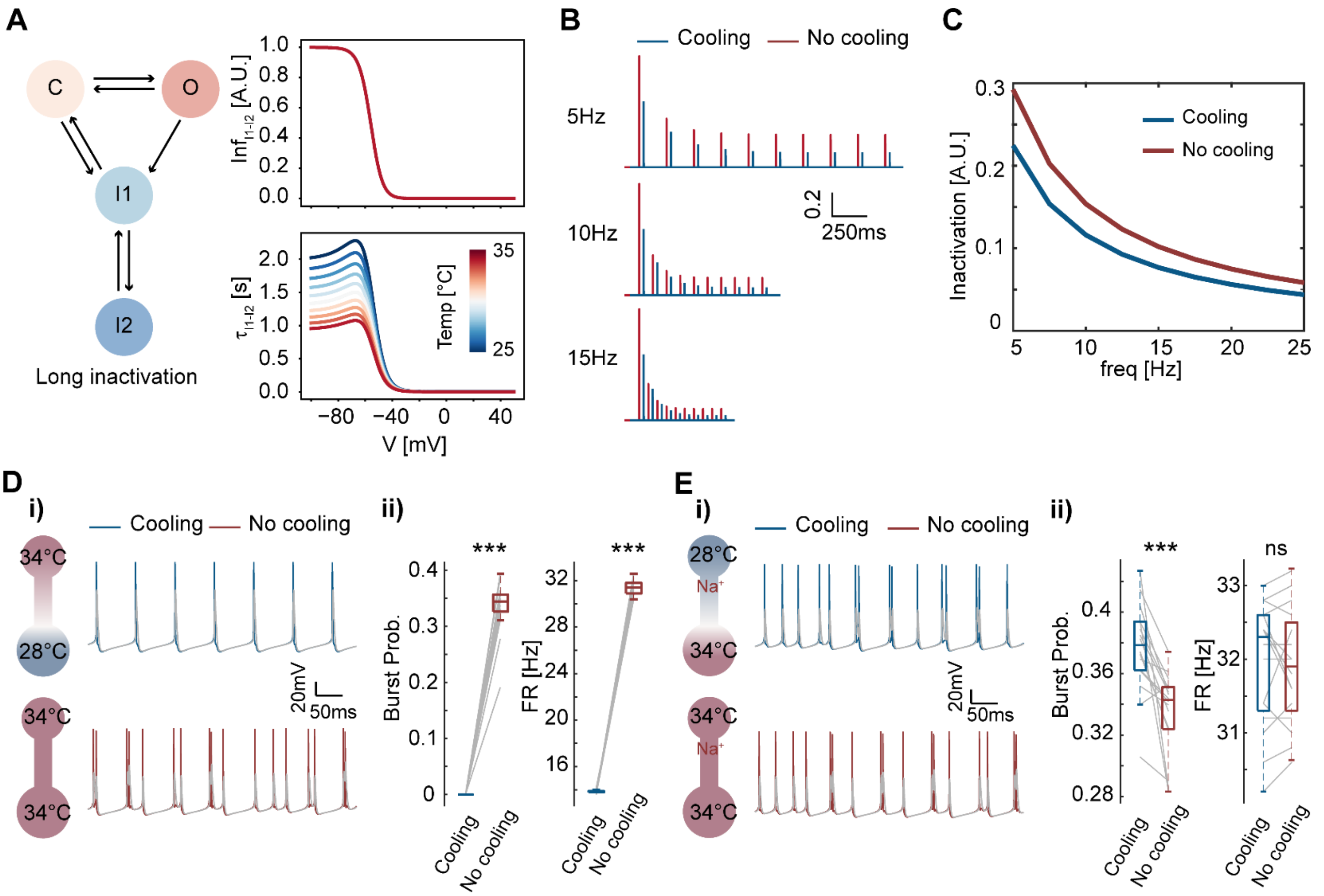


***Fig. S8. Slow recovery of Na⁺ channels from long-term inactivation resulted from cooling effects burstiness***

***A)*** *(Left) Gating kinetic of the dendritic Na⁺ channels. (Right) Steady-state (top) and time constant (bottom) of the long-term inactivation gate ($I_2$) as a function of temperature.* ***B)*** *Dendritic Na⁺ channel gating under 10 short voltage pulses at different frequencies under mild cooling (blue, 28°C) and physiological temperature (red, 34°C).* ***C)*** *Percentage of Na⁺ channels opening at the $10^{th}$ pulse compared to the $1^{st}$ pulse under cooling (blue) and physiological temperature (red).* ***D)*** *i) Simulated spiking activities under somatic focal cooling (top) and physiological temperature (bottom), with gray traces indicating corresponding dendritic voltage. ii) Both burst probability and firing rate are reduced by somatic focal cooling (n = 20, Wilcoxon signed rank test, ***$p < 0.001$).* ***E)*** Similar to D), but with focal cooling on apical dendrites, wherein dendritic Na⁺ channels exhibit no temperature modulation (n = 16, Wilcoxon signed rank test, ***$p < 0.001$).

**Table S1: Parameters for the biophysical model**

| Compartment | Parameter name | Value |
|---|---|---|
| global | e_na | 50 mV |
| | e_k | -90 mV |
| | e_ca | 140 mV |
| | membrane time constant | 25 ms |
| | Ca2+ diffuse gamma | 0.0006 |
| | Ca2+ diffuse time constant | 35.7 ms |
| soma-AIS | g_leak | 0.03 mS/cm^2 |
| | Cm | 0.75 µf/cm^2 |
| | g_na | 3000 mS/cm^2 |
| | g_k | 300 mS/cm^2 |
| | area | 600 µm^2 |
| Distal apical | g_leak | 0.03 mS/cm^2 |
| | Cm | 0.75 µf/cm^2 |
| | g_nad | 20 mS/cm^2 |
| | Im | 0.1 mS/cm^2 |
| | g_ca | 0.3 mS/cm^2 |
| | g_kca | 3 mS/cm^2 |
| | distance to soma | 400 µm |
| | rho (ratio of area to somatic area) | 20 |
| | kappa (axial coupling resistance) | 5 Mohm |
| Proximal apical | g_leak | 0.03 mS/cm^2 |
| | Cm | 0.75 µf/cm^2 |
| | g_nad | 20 mS/cm^2 |
| | g_k | 0.4 mS/cm^2 |
| | distance to soma | 250 µm |
| | rho (ratio of area to somatic area) | 15 |
| | kappa (axial coupling resistance) | 5 Mohm |
